# Habitat amount, temperature and biotic interactions drive community structure, life-history traits, and performance traits of cavity-nesting bees and wasps and their natural enemies in cities

**DOI:** 10.1101/2023.01.07.522464

**Authors:** Joan Casanelles-Abella, Loïc Pellissier, Cristiana Aleixo, Marta Alós Orti, François Chiron, Nicolas Deguines, Lauri Laanisto, Łukasz Myczko, Ülo Niinemets, Pedro Pinho, Roeland Samson, Piotr Tryjanowski, Lucía Villarroya-Villalba, Marco Moretti

## Abstract

1. Urban ecosystems are associated with socio-ecological conditions that can filter and promote taxa. However, the strength of the effect of ecological filtering on biodiversity could vary among biotic and abiotic factors. Here, we investigate the effects of habitat amount, temperature, and host-enemy biotic interactions in shaping communities of cavity-nesting bees and wasps (CNBW) and their natural enemies.
2. We installed trap-nests in 80 sites distributed along urban intensity gradients in 5 European cities (Antwerp, Paris, Poznan, Tartu and Zurich). We quantified the species richness and abundance of CNBW hosts and their natural enemies, as well as two performance traits (survival and parasitism) and two life-history traits (sex ratio and number of offspring per nest for the hosts). We analysed the importance of the abiotic and biotic variables using generalized linear models and multi-model inference.
3. We found that habitat amount and temperature were the main drivers of CNBW host responses, with larger habitat amounts resulting in higher species richness and abundance, and a larger total number of brood cells per nest for both bees and wasps, as well as a larger probability of survival for bees. Conversely, higher local temperatures decreased species richness, abundance, survival rate, number of brood cells per nest, and proportion of females in CNBW hosts.
4. Biotic interactions with natural enemies shaped wasp species richness, with higher levels of parasitism resulting in more wasp species. Similarly, our results showed direct density-dependence between CNBW hosts and their natural enemies.
5. Overall, our study highlights the importance of habitat amount and temperature in shaping urban food webs, through direct effects on hosts responses and the subsequent consequences for their natural enemies. As cities prepare to tackle the future consequences of global change, strategies that make it possible to maintain available habitat and mitigate urban overheating emerge as a key urban adaptation for biodiversity conservation.

## 1. INTRODUCTION

Urbanization both filters and promotes biodiversity, providing challenges and opportunities for urban wildlife management. Understanding the drivers of urban diversity patterns and ecological processes across urban ecosystems has therefore become a central part of urban ecology research to promote biodiverse cities (Uchida et al., 2021). Urban diversity patterns are structured following pronounced socio-ecological gradients (Uchida et al., 2021), i.e. urban intensity gradients, which structure species assemblages (Casanelles-Abella, Chauvier et al., 2021). Nonetheless, how urbanization affects multitrophic biodiversity characteristics at lower organizational levels, such as performance traits (e.g. parasitism, mortality, survival; *sensu* Violle et al., 2007) and life-history traits (e.g. sex ratio, number of eggs), could help unravel species fitness, survival and occupancy responses, allowing more detailed management actions. Species assemblages and the fitness of single species are determined by environmental conditions, such as habitat amount (i.e. the total area of suitable habitats at a defined space; *sensu* Fahrig, 2013), including food resource availability, and temperature, in combination with biotic interactions, such as competition for existing resources (Martin & Bonier, 2018), top-down control by higher trophic levels (Dainese et al., 2018), and bottom-up control by lower trophic levels (Steffan-Dewenter & Schiele, 2008).

Habitat amount has been identified as a main driver of ecosystem biodiversity (Fahrig, 2013; Hutchinson, 1959; Srivastava & Lawton, 1998), but the relationship between habitat amount and biodiversity might be more complex in cities than in other ecosystems. Species richness generally increases when the amount of habitat within an appropriately defined area is larger (Fahrig, 2013), due to the combined effects of complementation and supplementation of resources (Colding, 2007; Dunning et al., 1992;). In addition, more diverse plant communities increase habitat heterogeneity, which cascade up to higher trophic levels (Fabian et al., 2014; Hutchinson, 1959). Greater plant diversity often results in more food resources (e.g. nectar), enhancing the abundance and richness of consumers directly, and of higher trophic levels indirectly (Fabian et al., 2014; Srivastava & Lawton, 1998). Besides affecting community-level responses, habitat amount also influences life-history traits. For instance, in many hymenoptera species, female offspring require a larger investment of food resources than male offspring, which might result in male-biased sex ratios when the amount of habitat and therefore food resources decrease (Fitch et al., 2019). The dependence on habitat amount is expected to vary according to the trophic level of the studied taxa and their mobility (Fabian et al., 2014; Mayr et al., 2020; Schuldt et al., 2019), with prior studies having mixed results (Albrecht et al., 2007; Ebeling et al., 2012; Fabian et al., 2014). Compared with semi-natural ecosystems, cities are characterized by small habitat patches subjected to variable levels of human interference, resulting in urban habitats that vary substantially in their properties (Aronson et al., 2017). Moreover, in cities, habitat amount and the distribution of complementary resources can also influence biotic interactions, such as antagonistic interactions, and therefore biotic controls, by promoting or limiting the dispersal of one or both interacting species (Burks & Philpott, 2017; da Rocha-Filho et al., 2020; Rocha & Fellowes, 2018). Consequently, how habitat amount influences the life-history traits, performance traits and community composition of urban species is less known than in other ecosystems but of major importance for biodiversity management in urban areas.

While temperature gradients and their effects on biodiversity are well documented in communities outside cities (Mayr et al., 2020; Orr et al., 2021), temperature changes across space have been also reported within cities (Steward & Oke, 2012). In particular, cities can have pronounced local temperature gradients, due to interactions between the atmosphere and the complex heterogeneity of the urban fabric. Different densities and impermeable materials and the presence of blue-green spaces can form urban heat or cooling islands at the points of greatest or fewest accumulation of heat and energy, respectively. High temperatures are expected to particularly favour ectothermic taxa (e.g. Banaszak-Cibicka et al., 2014; Geppert et al., 2022). However, excessively high temperatures, above a species’ physiological threshold at different life stages, could enhance desiccation and reduce survival and reproduction success (Dale & Frank, 2018), as well as creating a phenological mismatch with the plant hosts (Papanikolaou et al., 2017). Further, because of the positive association between temperature and grey surfaces, warmer areas can correspond to smaller amounts of green habitat and less availability of food resources. The effects of broad temperature gradients on biodiversity have been studied previously (e.g. Trøjelsgaard & Olesen, 2012), yet local temperature effects, particularly in cities, have been less investigated.

Biotic interactions between hosts and their natural enemies are an additional driver of diversity. Natural enemies (also known as natural antagonists) include predators and parasites, which can indirectly favour species richness by alleviating competitive pressure by dominant species (i.e. top-down regulation; Steffan-Dewenter & Schiele, 2008; Levi et al., 2019). Focusing on parasitism, interactions between hosts and their natural enemies can be either directly density-dependent (Dainese et al., 2018), when larger numbers of hosts result in increased parasitism (Hassell, 2000), or inversely density-dependent, when larger numbers of hosts result in decreased parasitism (Rosenheim, 1990; Steffan-Dewenter & Schiele, 2008). The dynamics between hosts and their natural enemies might be altered in cities, as the urban environment can affect the behaviour, distribution, physiology and community structure of both hosts and natural enemies, due to the combined action of multiple socio-ecological factors and processes (Classen-Rodríguez, et al., 202). However, studies on the effects of biotic interactions in cities are still scarce (Theodorou, 2022).

Cavity-nesting bees and wasps (CNBW) and their natural enemies offer a highly standard and replicable model system to test for the effects of potential abiotic and biotic drivers on multitrophic biodiversity and food webs. CNBW represent an ensemble of trophic levels including pollinators (bees), predators on different insect groups (wasps), and natural enemies (parasites, cleptoparasites, parasitoids) (Staab et al., 2018). CNBW and their natural enemies are particularly interesting to monitor within urban ecosystems, as they represent bioindicators for habitat quality and environmental change, yet they remain far less studied in this setting (but see Moretti et al., 2021; Pereira-Peixoto et al., 2014; Xie et al., 2022).

Here, we studied the effects of temperature, habitat amount, and biotic interactions between the hosts and their natural enemies on multitrophic diversity. In particular we considered effects on the community structure (i.e. species richness, abundance) of CNBW and their natural enemies, as well as on the performance (i.e. parasitism, survival) and life-history traits (i.e. sex ratio, number of offspring cells per nest) of CNBW across urban ecosystems. We tested the following non-mutually exclusive hypotheses:

1. Small amounts of habitat (i.e. when urban intensity is high), and consequently low food resource availability, reduce species richness and abundance of CNBW, the number of offspring per nest, the offspring survival, and reduce the number of females. This effect might be stronger in wasps than in bees, as loss of habitat particularly affects arthropod prey diversity and abundance at higher trophic levels (Attwood et al., 2008; Mayr et al., 2020). Similarly, smaller habitat amounts negatively impact natural enemies, ultimately decreasing parasitism rates.
2. Higher temperature, within physiological thresholds, positively influences the foraging activity of CNBW, resulting in higher species richness, abundance, number of eggs laid per nest, offspring survival, and the number of females (Geppert et al., 2022; Mayr et al., 2020).
3. Biotic interactions (host–natural-enemy interactions) might positively shape both hosts and natural enemies. Regarding hosts, higher parasitism rates and a larger number of parasitized brood cells might positively regulate host populations of bees and wasps by reducing competition among dominant species, thus increasing species richness. Regarding natural enemies, increased host availability (i.e. more nests and higher host species richness) positively influences the species richness of natural enemies and the number of parasitized cells by providing more resources (Dainese et al., 2018; Hassell, 1982; Rosenheim, 1990).
4. Conversely, biotic interactions might negatively influence both hosts and natural enemies. Regarding hosts, higher parasitism rates and a larger number of parasitized brood cells might reduce host population sizes. Regarding natural enemies, increased host availability (i.e. more nests and higher host species richness) might result in a greater probability of detection of natural enemies by the host, less time for natural enemies to lay eggs, and enhanced collective defences by hosts, ultimately reducing natural-enemy species richness and the number of parasitized cells (Rosenheim, 1990).

## 2. MATERIALS AND METHODS

### 2.1. Study cities and sites

Our study was set in five European cities: Antwerp (Belgium), Greater Paris (France, hereafter referred to as Paris), Poznan (Poland), Tartu (Estonia), and Zurich (Switzerland), covering a large part of the climatic variability in mainland Europe (Figure 1). Site selection was done using an orthogonal gradient of patch size and isolation (Text S1). Overall, the final selection included 80 sites, 32 in Zurich and 12 in each of the remaining four cities (Figure 1; Table S1). Sites were at least 500 m apart from each other, except for two sites in Zurich that were 260 m apart (see Text S1 for additional details).

**FIGURE 1.**
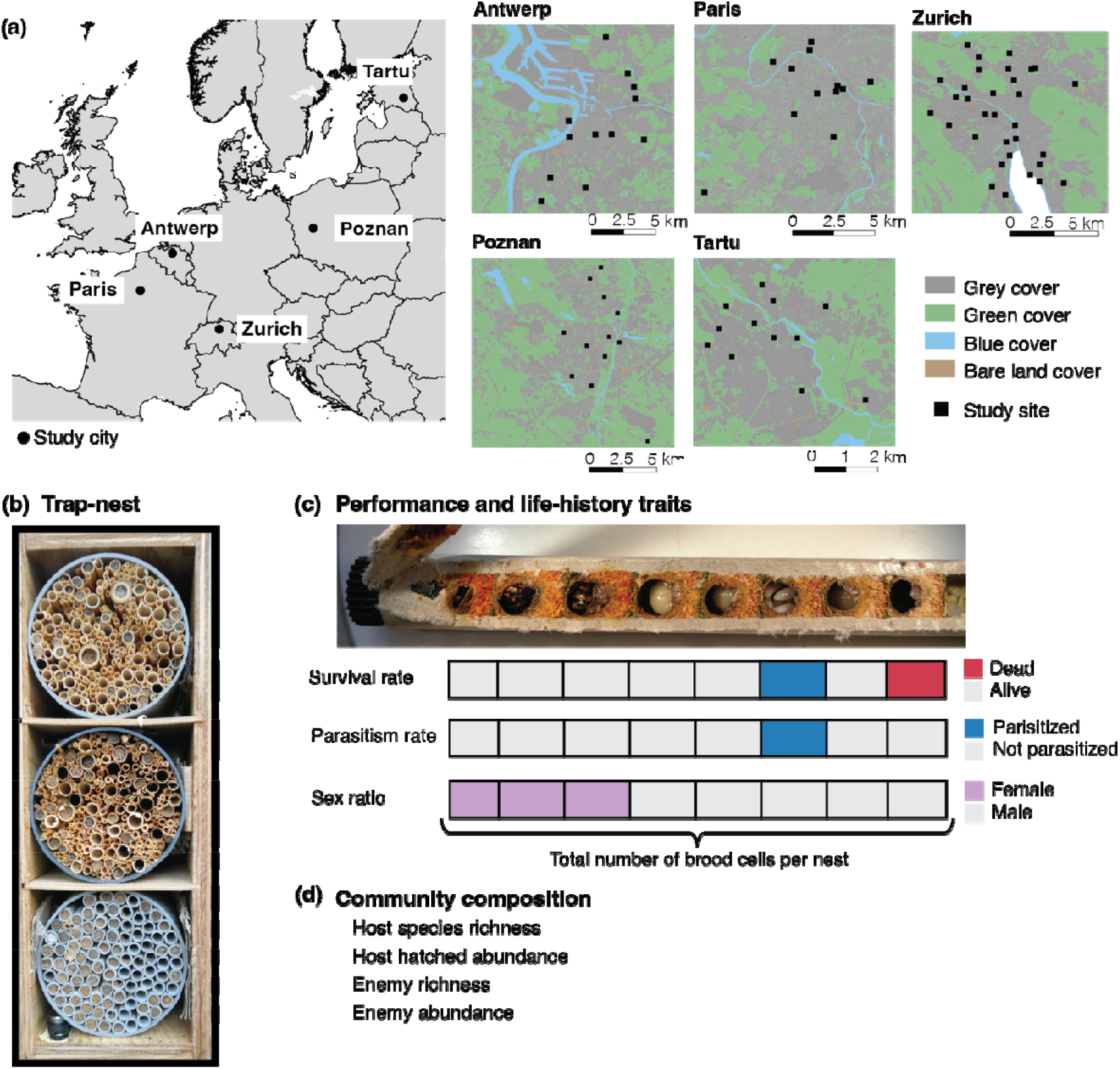
Study design and calculation of the response variables. (a) Study cities (Antwerp, Paris, Poznan, Tartu and Zurich) and distribution of study sites per city. (b) One of the trap-nests used in the study site Bois de Vincennes, Paris. (c) Calculation of ecological rates (survival rate and parasitism rate) and life-history traits (sex ratio and number of cells per nest) using a nest from *Osmia leaiana* (Kirby, 1802) in Zurich as an example. For each brood cell, information on survival, parasitism and sex was recorded in binary form (0 = dead, non-parasitized, male; 1 = alive, parasitized, female). The survival rate is the number of brood cells with emerging hosts, thus excluding the cells where the host died due to parasitism or other causes, divided by the total number of brood cells per nest. The parasitism rate is the number of brood cells where parasites were present divided by the total number of brood cells per nest. The sex ratio is expressed in terms of the proportion of females. Each rectangle represents an individual cell in the nest. (d) Selected community composition metrics for both hosts and natural enemies. Note that each metric was calculated separately for bees and wasps.

### 2.2. Insect sampling

#### 2.2.1. Trap-nests

We sampled cavity-nesting bees and wasps (CNBW) and their natural enemies using trap-nests. This enabled us to study their individual- and community-level responses in standardized nesting conditions (Figure 1b, see Text S2 for more details; Staab et al., 2018). To avoid vandalism, we installed trap-nests at 2.5–3.5 m height in sun-exposed SE or SW expositions for the period of January to October 2018. In October, we collected the trap-nests and stored them at c. 5°C. Between November and December 2018, we opened all reed internodes to detect brood cells. We counted the number of brood cells in the bee and wasp nests and noted the occurrence of natural enemies (cleptoparasites, parasites, parasitoids and predators). We then placed the reeds individually into glass tubes and closed the tubes with cotton wadding to collect emerging insects. We stored the tubes at 5°C until February 2019, when we transferred the tubes to a room at ambient temperature.

During spring and summer 2019, bees, wasps and their natural enemies emerged. We identified bees, wasps and enemies to the species level or to the lowest taxonomic rank possible (Table S2), using existing identification keys (Text S2). In addition, we identified the sex of all emerged bees and wasps and calculated the sex ratio at the nest level (i.e. for each reed tube). When no adult hosts emerged from a nest, we used the nest features (sealing material, diameter, larval food type) to identify the host genus or the family. Finally, some natural enemy species could only be identified to morphospecies (Text S1, Table S3).

#### 2.2.2. Community composition, performance traits and life-history traits

We calculated community composition, performance traits, and life-history traits (*sensu* Violle et al., 2007). Concerning the community structure, at each site we calculated species richness, abundance, and number of nests of the hatched hosts separately for bees and wasps. We calculated the abundance of hatched hosts as the number of brood cells where an individual emerged, i.e. the total number of brood cells minus the number of parasitized cells and the number of cells with no emergence for reasons other than parasitism. We calculated the number of nests as the total number of colonized reeds. The abundance of hatched hosts and number of nests were highly correlated (Pearson correlations from 0.81 to 0.92). Thus, we only included the abundance of hatched hosts in the analyses, as it reflects the actual number of individuals that will constitute the next generation. For natural enemies, we calculated the species richness and abundance. We estimated the abundance of natural enemies as the number of parasitized brood cells rather than the total number of emerging individual enemies, to account for differences in the number of eggs laid per cell across different enemy species.

We assessed performance traits, i.e. survival and parasitism, at the individual cell level. Specifically, for each cell within a nest, we noted whether the cell was alive or dead (survival; Figure 1c), and whether it had a parasite or not (parasitism; Figure 1c). Further, we calculated the parasitism rate as the number of cells with a parasite divided by the total number of cells. Concerning life-history traits, we calculated the total number of brood cells per nest as the total number of brood cells within a single nest, regardless of whether the larvae were dead, alive or parasitized (Figure 1c). Finally, we used the probability that a femalehost emerged from a given cell as a proxy for the sex ratio. To do so, we first identified the sex of the individual hosts. Then, we noted it in a binary way to indicate whether the host was a female or not (Figure 1c). We could identify the sex of all hatched hosts. Further, because CNBW first lay females and then males, on some occasions we could also identify the host sex in dead and parasitized cells, that is, when we had already identified the sex of the preceding and the following cells.

### 2.3. Habitat amount, temperature, and biotic interactions

We inferred habitat amount using a combination of landcover data and remote-sensing-based indices (Table S4). First, we estimated habitat amount using metrics based on land-cover data. Using the European Urban Atlas (EEA, 2010), we calculated the local patch size (total area of the study site) and patch isolation (using the proximity index, which weights the contribution of the area of neighbouring patches by the distance to the focal patch within a given search radius). We also assessed habitat amount by using the local landcover map developed by Alós-Ortí et al. (2022). This local landcover map distinguishes between vegetation and build-up types (grasslands, woody vegetation, and impervious surfaces within 32 m) and can be used an indirect proxy for habitat amount and particularly for food resource availability for bees and wasps.

Further, following the *habitat amount hypothesis* (Fahrig, 2013), which defines habitat amount as the total area of habitat at a defined distance from the nest, we used remote-sensing-based indices, calculated using 50, 100, 200, 400 and 800 m buffers from the focal trap-nests, to infer the habitat amount at each site. The selected buffer sizes are expected to be relevant for the studied organisms. In particular, we estimated the habitat amount using the normalized difference vegetation index (NDVI) within the five buffers. NDVI can be used to characterize existing vegetation and urban infrastructure and is related to the habitat suitability of species (Text S4). Due to the correlations between buffers, we retained only two scales in the models: 100 m and 800 m. Finally, we used the floristic inventories from Casanelles-Abella, Frey et al. (2021) to further characterize habitat amount in terms of food resource availability, inferred as plant species richness. Briefly, we performed the floristic inventories in standardized plots, documenting the occurrence of all entomophilous plants (excluding families Poaceae and Cyperaceae) within a 100 m buffer of the trap-nests. Floristic inventories took place on four occasions between April and July 2018.

We measured temperature at the trap-nest level and within a 800 m buffer from the trap-nests. At the trap-nest level, data loggers (1-Wire/Data Logger model DS1921G-F5, Analog Devices, Wilmington, MA, USA) recorded temperature hourly between February and June 2018, and we calculated the mean temperature per study site over this period (local temperature). We expect this scale of measurement to reflect the local thermal nesting conditions. Within the 800 m buffer (landscape temperature), we inferred temperature using remote sensing indices of land surface temperature (LST; Text S4). We expect this scale of measurement to reflect the thermal landscape surrounding our study sites. Additionally, we calculated LST for 50, 100, 200 and 400 m buffers, but ultimately did not use these estimates due to high inter-correlations (Pearson *r* > 0.8). The distribution of values of the used predictors can be found in Figure S1.

Finally, regarding biotic interactions, we used the number of parasitized cells and the species richness of natural enemies as top-down proxies (Table S4). For bottom-up proxies, we used the number of nests and the species richness of hosts (Table S4).

### 2.4. Statistical analyses

We used R version 4.0.2 (R Core Team, 2022) and RStudio v.07.1 (RStudio Team, 2020) for all analyses and statistical figures, using the packages glmmTMB v.1.1.3 (Brooks et al., 2017), MuMIn v.1.46.0 (Barton & Šímová, 2015), evaluate v.015 (H & Y, 2022), performance v.0.9.2 (Lüdecke et al., 2021), DHARMa v.0.4.5 (Hartig, 2022), and ggplot2 v.3.3.6 (Wickham, 2016).

We used generalized linear mixed-effects models (GLMMs) to assess the influence of temperature, habitat amount, and host–natural-enemy interactions (biotic interactions) on host and natural-enemy responses. For hosts, we considered the response variables species richness, abundance hatched, survival (probability that the host in a cell survived), parasitism (probability that the host in a cell was parasitized), sex ratio (probability that the host in a cell was female) and number of brood cells per nest. For natural enemies, we considered the response variables species richness and number of parasitized cells (proxy for enemy abundance). We ran models separately for bees and wasps and for the natural enemies of bees and wasps. We modelled species richness and abundance responses with a Poisson error structure when there was no overdispersion, and a negative binomial error structure and a log-link when we detected overdispersion. We built the models using city as a random term and a variable number of fixed effects (see below). For host parasitism, host survival, and host sex-ratio, we used cell-level data, encoding each variable as a binary output (1 = non-parasitized/alive/female; 0 = parasitized/dead/male). We used a binomial distribution with a logit-link (Zuur et al., 2010), using a nested random term (individual nest within site within city) and a variable number of fixed effects. For the number of brood cells per nest we used a negative binomial error structure, as overdispersion was detected, using a nested random term (site within city) and a variable number of fixed effects. One site in Antwerp (An057) was not colonized and was excluded from the analyses, leading to a sample size of 79 sites. We checked collinearity among the predictors using the variance inflation factor (VIF) and the Pearson correlation coefficient. We discarded variables with VIF > 3 and a Pearson correlation coefficient > 0.7 (Dormann et al., 2013; Zuur et al., 2010). Prior to the analyses, we standardized all predictors by z-transformation.

We performed multimodel inference to study CNBW responses. We first formulated a global mixed model that included all fixed effects and the random structure using the R package glmmTMB. We used diagnostic plots to estimate model performance and test for spatial autocorrelation. Then, we ran all possible combinations of nested models with the function ‘dredge’ in the R package MuMIn to automatically construct all possible models based on the set of predictors. For model evaluation and selection, we used the Akaike information criterion corrected for small sample sizes (AICc; Burnham et al., 2011) to select the best model. If more than one plausible model existed (i.e. Δ AICc < 6, following Burnham et al., 2011 and Magrach et al., 2017), to account for model selection uncertainty we proceeded to compute the full-model averaged parameter estimates for each predicting variable in the candidate model set, using zero when predictors were not included in a particular model (Symonds and Moussali, 2011). The relative importance of a predictor was based on the sum of the Akaike weights across all models in the candidate model set that included the predictor (Burnham et al., 2011).

## 3. RESULTS

We found a total of 4392 nests of cavity-nesting bees (1998 nests) and wasps (2394 nests), containing 16,617 brood cells from 16 bee and 45 wasp species (Table S5), and 4 bee and 14 wasp morphospecies (Figure S2). Four species (*Chelostoma florisomne, Hylaeus communis, Osmia bicornis* and *Osmia cornuta*) accounted for about 80% of the total number of bee brood cells (Table S2), whereas for wasps the total number of brood cells was more evenly distributed across species, indicating a greater dominance in cavity-nesting bees. All species were native, with the exception of the sphecid wasp *Isodontia mexicana* (Saussure, 1867), which was recorded in Antwerp, Paris and Zurich. Further, we identified a total of 47 natural enemy species or morphospecies in 812 nests and 1500 brood cells, representing taxa from three insect orders (Hymenoptera, Coleoptera, Diptera) and mites (*Chaetodactylus* spp.).

### 3.1. Effect of habitat amount

We found host community structure (species richness and abundance hatched) to be strongly associated with the proxies for habitat amount (i.e. NDVI value within a given buffer from the trap-nest) for both bees and wasps. In particular, sites with higher NDVI values had higher host species richness and abundance of the hatched hosts (Tables 1 and S6, Figure 2a,b). More classical predictors representing habitat amount, i.e. patch area and proximity index (proxy for isolation), were never significant (Tables 1 and S6, Figure S3), although patch area was positively correlated with both species richness and abundance of CNBW (Figure 2d,e). On the other hand, habitat amount did not directly influence natural enemies (Tables 1 and S6, Figure S4).

**TABLE 1.**
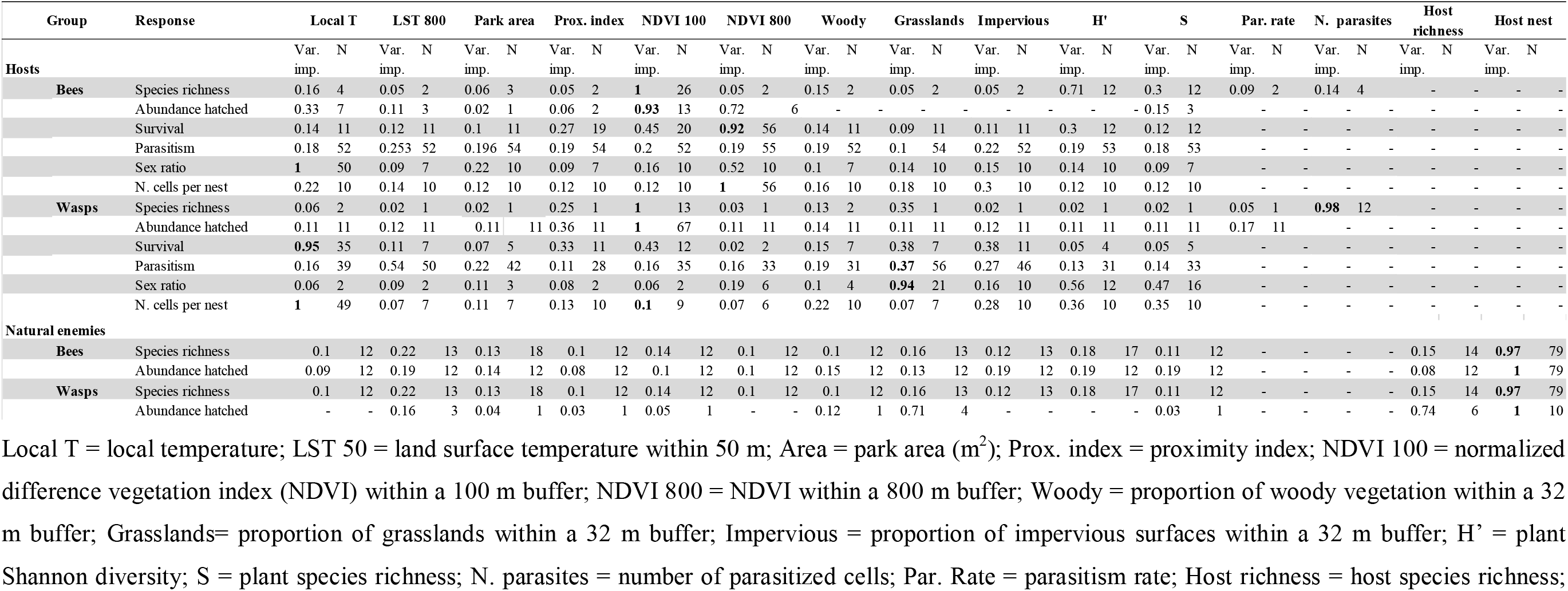
Relative importance (Var. imp.) of the potential predictors in determining the responses of cavity-nesting bees and wasps and their natural enemies after model averaging. The number of models (Δ AICc < 6) containing a particular variable is also given (N). Statistically significant variables (p < 0.05) are given in bold. Variables with a relative importance > 0.7 were considered good predictors for the target variable.

**FIGURE 2.**
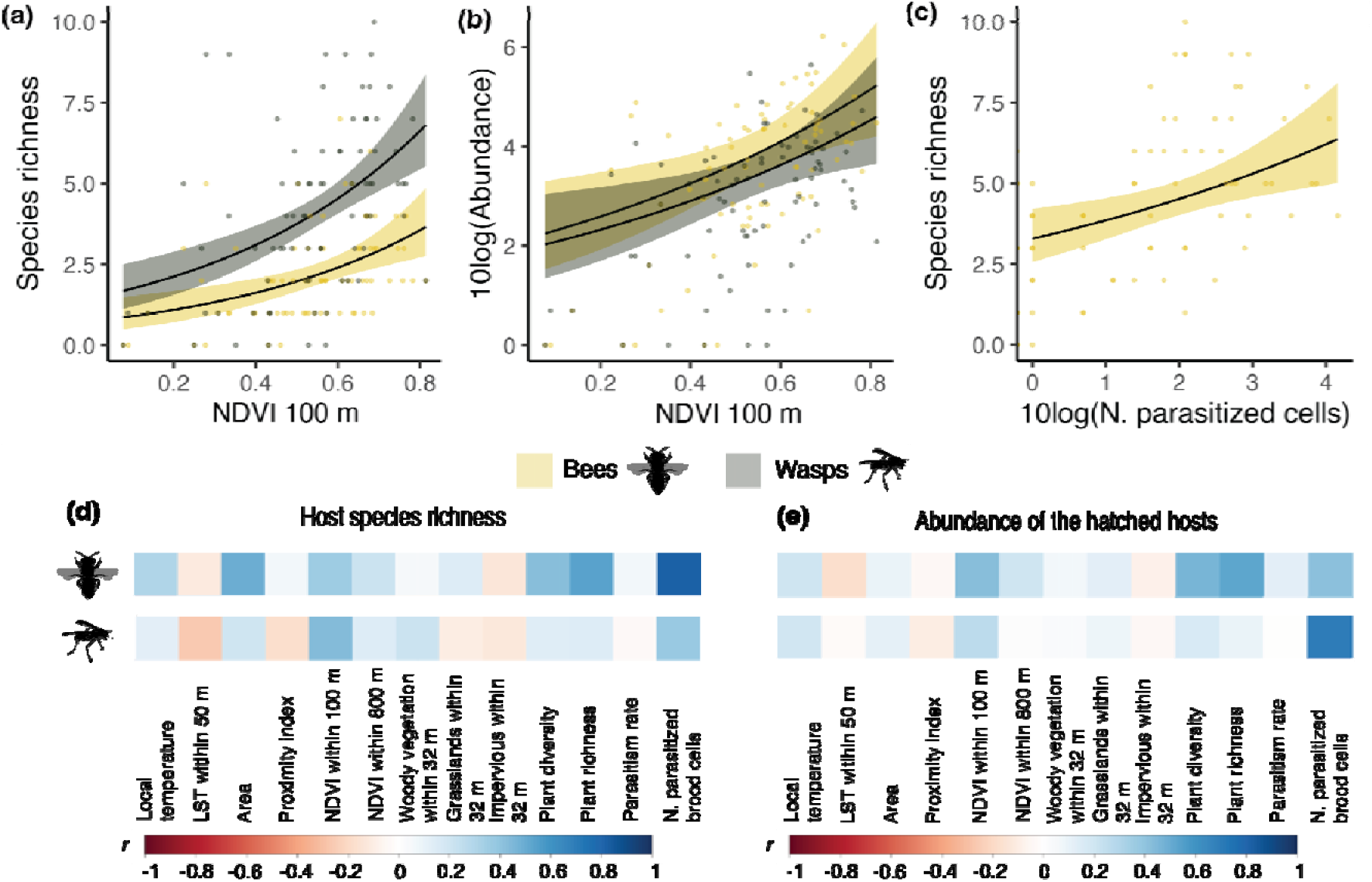
Drivers of the community composition of cavity-nesting bees and wasps. (a–d) Generalized linear model (GLMM) results for the relationship between the predicted metrics of community composition (species richness and abundance hatched) of the cavity-nesting bees and wasps plotted against proxies for (a, b) habitat amount, represented by NDVI within a 100 m radius and (c) biotic interactions, represented by the number of parasitized brood cells. Points represent the observed data and solid lines represent the predicted values obtained using averaged GLMMs; coloured bands indicate 95% confidence intervals. A summary of the model results is provided in Table 1 and details are provided in Table S6. (d–e) Pearson correlations between (d) the species richness or (e) the abundance of cavity-nesting bees (top row) and wasps (bottom row) and the proxies for temperature, habitat amount and biotic interactions. Abundance is expressed as the total number of hatched offspring. For further relationships see Figure S3.

Multi-model inference revealed the importance of the different drivers in shaping the studied performance traits and life-history traits of CNBW. Habitat amount was the main driver of bee survival for the vast majority of models computed, with the probability of survival increasing in sites with higher NDVI values (Figure 4). Other proxies for habitat amount at smaller spatial scales, such as NDVI within 100 m and plant diversity within 50 m, were retained in several models and had a positive but not significant relationship with bee survival (Table S6, Figure S5). Finally, for bees the total number of cells per nest was mainly (positively) affected by increasing habitat amount at large spatial scales (represented by NDVI within 800 m), and this variable was retained in the vast majority of the models (Figure 4, Table S6).

**FIGURE 3.**
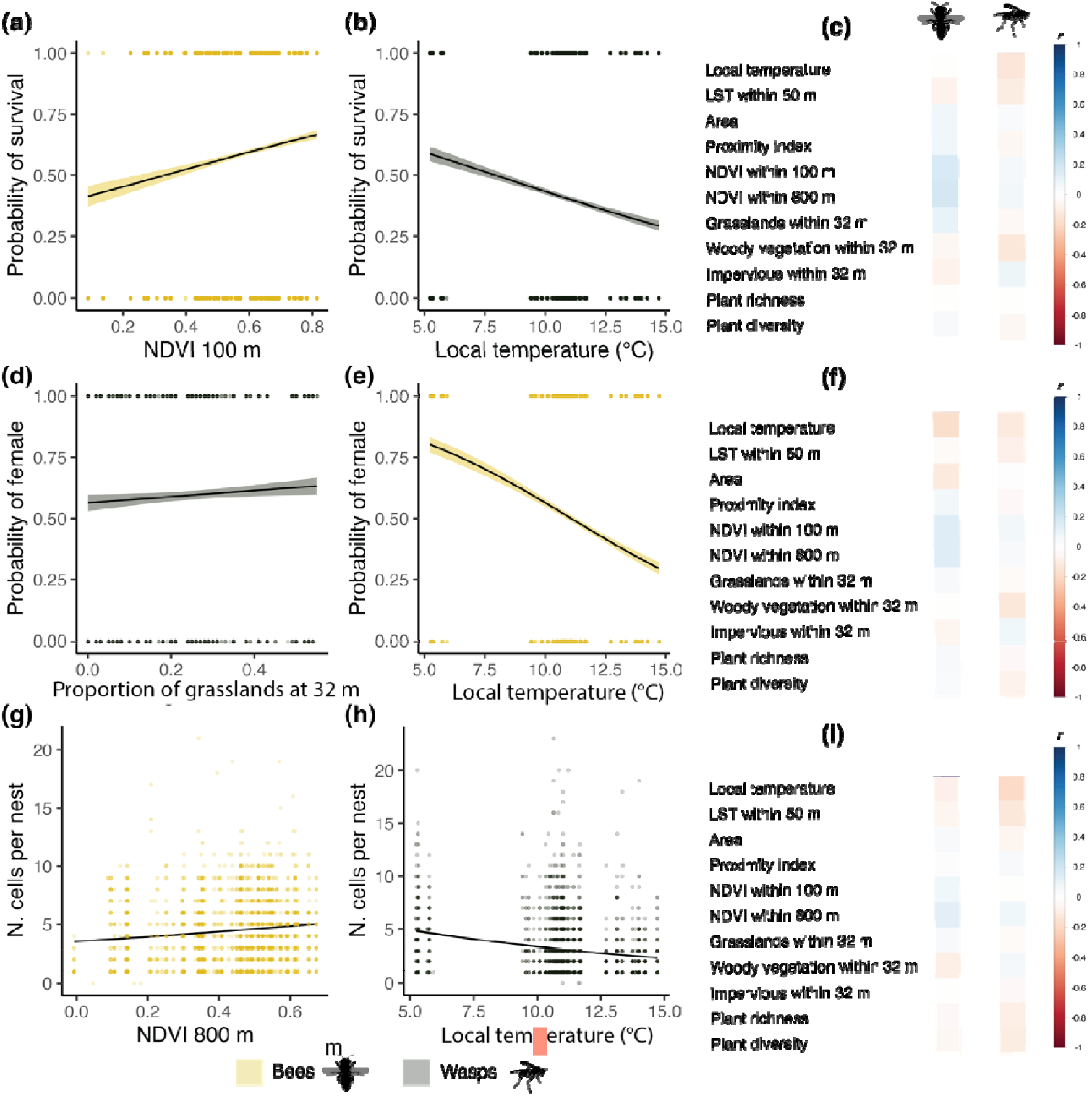
Drivers of the ecological rates and life-history traits of cavity-nesting bees and wasps. Generalized linear model (GLMM) results for relationships between the predicted metrics of performance traits (a–b: probability of survival, i.e. probability that the host in a cell survived), and life-history traits (d–e: probability of female sex, i.e. probability that the host in a cell was female; g–h: number of brood cells per nest) of cavity-nesting bees and wasps. Points represent the observed data and solid lines represent the predicted values obtained using averaged GLMMs; coloured bands indicate 95% confidence intervals. Model results are shown in Table S6. (c, f & i) Pearson correlations between the different predictors representing habitat amount and temperature and (c) survival rate, (f) proportion of females, and (i) number of brood cells per nest. The survival rate, parasitism rate, proportion of females, and number of brood cells per nest are calculated at the nest level. For further relationships see Figure S5.

**FIGURE 4.**
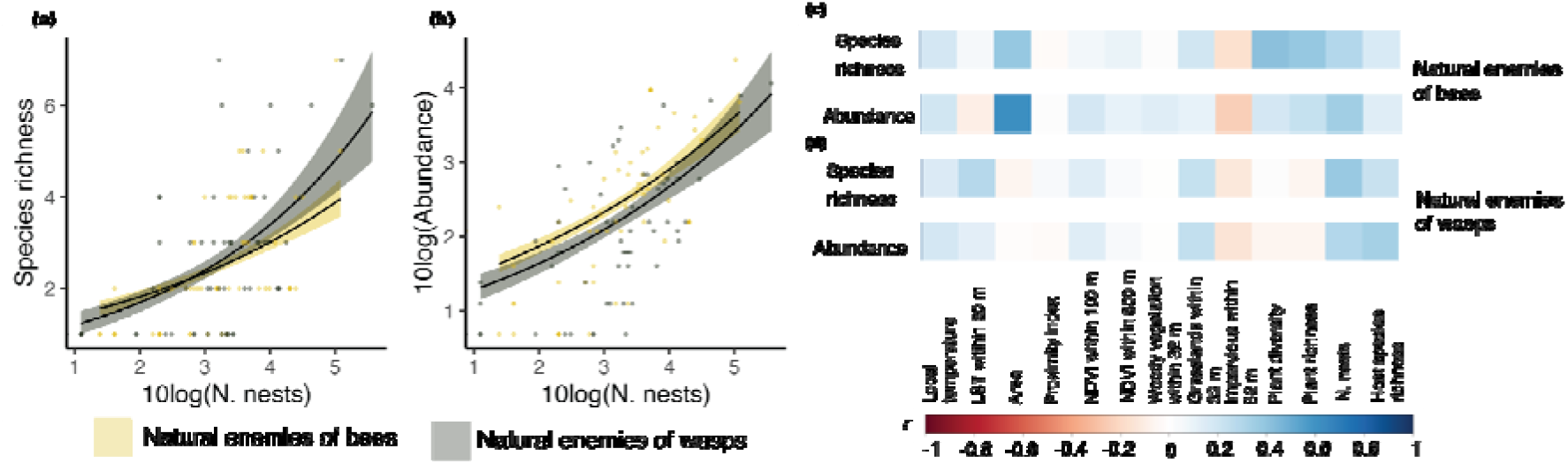
Drivers of the natural-enemy community. (a–b) Generalized linear model (GLMM) results for relationships between the predicted metrics of (a) species richness or (b) abundance (10log-transformed) of the natural enemies (yellow and grey represent natural enemies of bees and wasps, respectively) and the proxies for bottom-up control or resource availability for natural enemies of cavity-nesting bees and wasps. Abundance is estimated as the number of parasitized brood cells. Points represent the observed data and solid lines represent the predicted values obtained using averaged GLMMs; coloured bands indicate 95% confidence intervals. Model results are shown in Table S6. (c–d) Pearson correlations between the species richness and abundance of the natural enemies of (c) bees or (d) wasps and the proxies for temperature, habitat amount and biotic interactions. For further relationships see Figure S4.

### 3.2. Effect of temperature

Temperature was associated with host species richness and abundance, but the effect varied according to the spatial scale considered. In particular, temperatures measured at the landscape scale (LST within 800 m buffer) were negatively correlated with bee and wasp species richness and abundance (Figures 2 and S3). Conversely, local temperatures were positively correlated with the species richness and abundance of hatched offspring for both bees and wasps (Figures 2 and S3). However, the effects of local and landscape temperature were not significant (Tables 1 and S6).

Regarding performance traits and life-history traits, local temperature was negatively associated with the survival of wasp offspring (Table 1). Wasp survival also increased with increasing values for habitat amount proxies (NDVI within 100 m, proximity index, impervious surfaces within 32 m), which were retained in several models but were not significant after model averaging (Tables 1 and S6, Figure S5). Higher local temperatures reduced the probability of a female bee emerging from a given brood cell (Figure 3, Tables 1 and S6), leading to a larger proportion of males. For wasps, the probability that a female emerged from a given brood cell increased at sites with a larger proportion of grasslands (Figure 3). Local temperature was the main driver of the total number of cells per wasp nest (Table 1; Figure 3), with the total number of cells per nest declining with increasing temperature (Figure 3, Tables 1 and S6).

### 3.3. Effect of biotic interactions

Biotic interactions shaped the species richness of wasps (Figure 2, Table S6). Specifically, a larger number of parasites positively influenced wasp species richness. The effect was non-significant for bees (Figure 2c). Finally, both the abundance and the species richness of natural enemies of both bees and wasps increased with a larger number of nests per site (Figure 4a,b, Tables 1, 2 and S6).

**TABLE 2.**
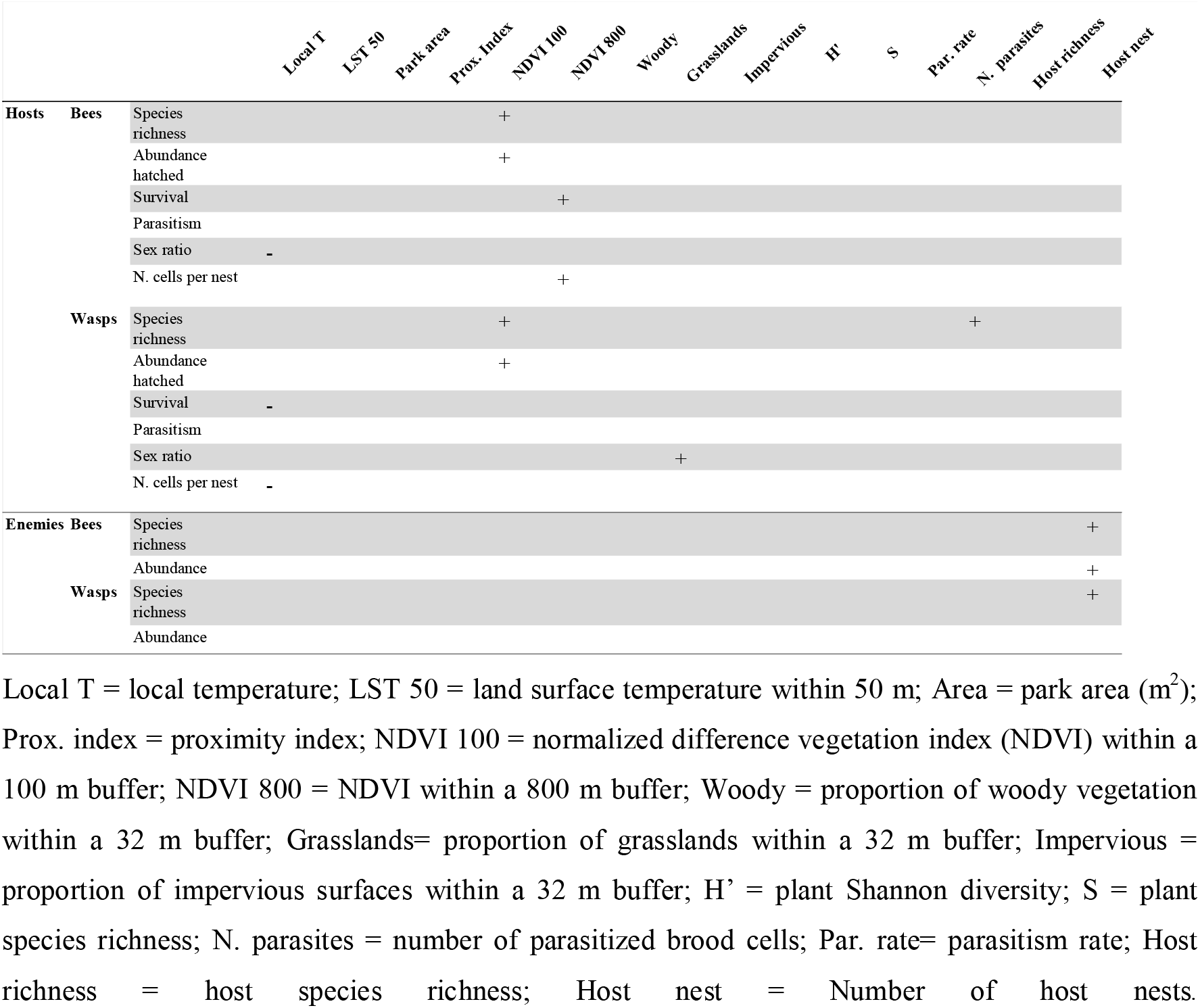
Summary of the effects of the proxies for temperature, habitat amount, resource availability and biotic controls from the full averaged models after performing multimodal inference on the responses of cavity-nesting bees and wasps and their natural enemies. Plus signs indicate a positive estimate whereas minus signs indicate a negative estimate (see Tables S6 for detailed information on the models’ input and Table S4 for additional information on the proxies).

## 4. DISCUSSION

Our results stress the importance of habitat amount and biotic interactions in enhancing multitrophic biodiversity in urban ecosystems. We show that the species richness, abundance and number of cells per nest of cavity-nesting bees and wasps (CNBW), as well as bee survival rates, increases in sites with higher NDVI values and therefore with smaller proportions of impervious surfaces and bare soil, as well as more vegetation cover. Greater vegetation cover has been found to be linked with higher abundances and richer communities of insect pollinators (Albrecht et al., 2009) and predators (Fabian et al., 2014). Moreover, our study supports the relevance of focusing on overall habitat amount, as proposed by Fahrig (2013), rather than on specific metrics of patch size and isolation. The *habitat amount hypothesis* (Fahrig, 2013) is very useful for understanding urban biodiversity patterns, as cities are characterized by a large number of relatively small, diverse, not necessarily adjacent patches (Aronson et al., 2017), explaining why certain cityscapes can still harbour relatively high diversity levels (Fournier et al., 2020; Vega & Küffer, 2021). Our study supports the idea that overall urban habitats, most of which are subjected to at least a certain degree of anthropogenic management (Aronson et al., 2016), contribute to maintaining different species assemblages across trophic levels and their interactions, due to habitat supplementation and complementation (Colding, 2007). This contrasts with what has been observed in regions or cities with shorter land-use-change histories (e.g. Brazil; da Rocha-Filho et al., 2019), where a smaller amount of natural habitat, rather than overall urban habitats, has been found to strongly negatively affect bee and wasp diversities.

The effects of habitat amount are complex and often indirect, and can vary across trophic levels (Mayr et al., 2020; Stefan-Dewenter & Schiele, 2008). In our study, habitat amount did not directly affect natural enemies. Instead, proxies for biotic interactions with their hosts, assessed using metrics representing the availability of hosts (i.e. number of nests, host species richness), were the main drivers, as observed in other studies (Dainese et al., 2018; Fabian et al., 2014). Nonetheless, the effects of habitat amount cannot be disregarded, as host communities depended strongly on the amount of habitat, hence indirectly affecting higher trophic levels. Indirect effects of habitat loss have been documented across food webs (Melián & Bascompte, 2003; Ryser et al., 2019), with negative effects occurring particularly for taxa from higher trophic levels (Melián & Bascompte, 2003).

Our metrics of plant diversity and vegetation composition, which we used as proxies for habitat amount and food resource availability, played only a minor role in shaping host responses, particularly for bees. Local food resource availability has been found to have a variable effect on the structure of CNBW communities in non-urban ecosystems (Albrecht et al., 2007; Mayr et al., 2020). On the one hand, plant and flower diversities have been observed to increase bee species richness and abundance in natural environments (Mayr et al., 2020) and restored natural meadows (Albrecht et al., 2007). On the other hand, no effect of plant or flower diversity on CNBW has been reported in other studies (Ebeling et al., 2012; Fabian et al., 2014), suggesting that the mobility and quantity of the food resource mediates the effects of plant diversity. Further advancements in the use of remotely sensed data to infer plant diversity and heterogeneity (e.g. Yang et al., 2022) at larger spatial scales could be a cost-effective solution to better test the specific effects of resource availability at the appropriate spatial scales. Further, vegetation metrics had a weak or null influence in our models, indicating that specific measurements of prey availability (e.g. abundance of spiders, aphids and caterpillars) could be a better proxy for food resource availability for wasps (Mayr et al., 2020).

While temperature can directly influence insect metabolism (Zuo et al., 2012) and can further regulate species richness and abundance, we did not find clear evidence of this in our studied cities. First, we found that higher temperatures at the local scale increased both species richness and abundance, while higher temperatures at the landscape scale had the opposite effect. Second, survival rate, the number of brood cells per nest, and the proportion of females tended to be reduced with increasing local temperature. Warmer environments have been found to enhance wild bee diversity at global (Orr et al., 2021) and city scales (e.g. in Rome, Geppert et al., 2022), while less is known about wasps. However, temperatures in cities are highly influenced by the amount and distribution of impervious surfaces, the density of buildings and the type of artificial materials. Therefore, higher temperatures, particularly when measured at landscape scales (e.g. LST within multiple buffer sized in our study), are also indicative of smaller habitat amounts and likely of lower food resource availability, which ultimately negatively impact biodiversity responses across ecological levels of organization, as we observed. For instance, in Zurich the species richness of wasps and of bees has been found to decline along urban intensity gradients and with lower vegetation heterogeneity and less urban green-space cover (Casanelles-Abella, Chauvier et al., 2021). In these circumstances, the effect of landscape temperature per se might be masked by the main effects of the loss of habitat and resources.

Another potential reason for the different effects of temperature depending on spatial scale of the measurements is that temperature at the trap-nest level was recorded only until June. This temporal range might have missed the species that nest later, although most CNBW build their nests between spring and early summer. Moreover, we did not measure the temperature during the months when the larval or pupal stages of many species are developing, i.e. from late summer to the following spring. Temperature sensitivity can vary during ontogeny (Rombough, 2003), and thermal conditions during the larval and pupal stages can therefore be critical for later CNBW development and emergence (Rombough, 2003; Ostap-Chec et al., 2021). Hence, future studies should extend temperature measurements to also include the all developmental stages of CNBW.

How biotic interactions influence the properties of host communities in cities remains poorly understood. Our results showed that wasp richness is positively regulated by their natural enemies, indicating a possible top-down regulation from natural enemies for wasps, whereas this effect was much weaker for bees. Thus, our results support the idea that natural enemies reduce competition, hence facilitating richer wasp species assemblages (Levi et al., 2019). This contrasts with findings from prior studies on cavity-nesting wasps in non-urban ecosystems, where resource availability (Steffan-Dewenter & Schiele, 2008) together with temperature (Mayr et al., 2020) were stronger drivers of CNBW diversity. Regarding bees, our finding of a lack of influence of top-down controls should be interpreted with caution, as other top-down factors, such as predation, were not assessed and could be more important (Vidal & Murphy, 2017). It is also possible that habitat, and subsequently resource availability, represent a greater limitation than parasitism pressure on bee hosts. For instance, a study on populations of the cavity-nesting bee *Osmia bicornis* in agricultural areas in Germany showed that resource availability was the primary driver, rather than regulation from natural enemies (Steffan-Dewenter & Schiele, 2008).

Biotic interactions were also the main driver of natural-enemy species richness and abundance, with a direct density-dependence on their hosts. Previous studies have also reported direct density-dependence (e.g. Krewenka et al., 2011; Mayr et al., 2020). Hence, our results suggest that the factors proposed to promote inversely density-dependent parasitism (i.e. limited handling time by natural enemies or improved defence and guarding capacities of aggregated hosts; see Rosenheim, 1990) do not apply to our study organisms. In fact, trap-nests force CNBW to nest in higher densities, which has often been observed to increase parasitism rates (Groulx & Forrest, 2018). Thus, CNBW, despite including gregarious species (i.e. *Osmia bicornis* and *Osmia cornuta;* Groulx & Forrest, 2018), do not appear to benefit from improved defences. Moreover, our findings underpin the dependence of natural enemies on their resources (hosts), supporting the idea that higher trophic levels are severely limited by their food resources (Mayr et al., 2020). In that regard, natural enemies are vulnerable to the decline in host availability and to the drivers responsible for these declines, such as habitat reduction, loss of key resources, and the spread of novel pathogens or diseases, which are well-known drivers of insect decline and are becoming more severe worldwide (Wagner et al., 2021).

## 5. CONCLUSIONS

Our study highlights the importance of habitat amount and temperature in urban food webs, through the direct effects on hosts responses and the subsequent consequences on their natural enemies. Ongoing climate change and its interaction with the urban fabric (Müller et al., 2014), together with urban intensification scenarios in the face of a growing urban population (Liu et al., 2020), therefore represent two main challenges for the survival of urban CNBW communities, as well as the interacting species in higher or lower trophic levels. Ongoing adaptation plans to reduce overheating through targeted greening and the expansion of novel and restored habitats represent an opportunity to maintain urban biodiversity (Butt et al., 2018), including the studied food webs. Overall, the study of multitrophic diversity improves our understanding of the contributions of different drivers to multiple dimensions of biodiversity, which in turn helps us to monitor ecological conditions and anticipate future challenges for biodiversity conservation.

## ACKNOWLEDGEMENTS

This research was supported by the Swiss National Science Foundation (project 31BD30_172467) within the ERA-Net Bio-divERsA project “BioVEINS: Connectivity of green and blue infrastructures: living veins for biodiverse and healthy cities” (H2020 BiodivERsA32015104) and by the Swiss Federal Office for the Environment (FOEN) in the framework of the project “City4Bees” (contract no. 16.0101.PJ/S284-0366). We also acknowledge Göhner Stiftung for supporting this project (2019-2917/1.1). CA was supported by the FCT (SFRH/BD/141822/2018). P.T. and Ł.M. acknowledge Polish funding through NCN/2016/22/Z/NZ8/00004. PP to FCT (CEECIND/03415/2020, DivRestore/001/2020). We thank the local authorities of Zurich (particularly Grün Stadt Zurich [GSZ]), Antwerp, Paris, Poznan and Tartu for supporting this study. Moreover, we acknowledge L. Roquer, J. Bosch and A. Rodrigo for their input on the trap-nest collection. We thank S. Müller, E. Eggenberg, K. Kilchhofer, P. Bischof, R. Veenstra, Ł. Myczko, A. Zanetta and D. Frey for help collecting and processing the bee data. We are grateful to S. Fontana and F. Duchenne for their guidance with the statistical analyses.

## Conflict of interest

The authors declare no conflict of interest.

## Notes

### Competing Interest Statement

The authors have declared no competing interest.

