## Supplementary material for "Habitat amount, temperature and biotic interactions drive community structure, life-history traits, and performance traits of cavity-nesting bees and wasps and their natural enemies in cities"

**Supplementary text:**
**Text S1.** Site selection.
**Text S2.** Insect sampling and identification.
**Text S3.** Details on local landcover map.
**Text S4.** Details on remote sensing metrics.

**Supplementary tables:**
**Table S1.** Study sites.
**Table S2.** List of cavity-nesting bee and wasp species.
**Table S3.** List of natural enemy species.
**Table S4.** Summary of variables.
**Table S5.** Community composition.
**Table S6.** Results from multimodel inference.

**Supplementary figures:**
**Figure S1.** Predictors across cities.
**Figure S2.** Community composition.
**Figure S3.** Species richness and abundance of hosts plotted against predictors.
**Figure S4.** Species richness and abundance of natural enemies plotted against predictors.
**Figure S5.** Performance traits and life-history traits plotted against predictors.

**TEXT S1. Site selection.** Urban green areas were selected using the pan-European Urban Atlas (EEA, 2012). Study sites were selected following an orthogonal gradient of local habitat amount (patch size) and landscape habitat amount (using the proximity index PI; McGarigal et al., 2012). Specifically, the proximity index weights the contribution of the area of neighbouring patches by the distance to the focal patch within a given search radius. To select the study sites, all possible sites were classified into six patch size classes and six classes of PI (36 possible combinations). Within these combinations, sites were selected randomly (random stratified sampling design). Due to resource limitations, only one-third of the possible combinations in Antwerp, Paris, Poznan and Tartu were used, whilst the full range of combinations was used in Zurich (32 combinations in total; the other combinations were not available). Further details can be found in Casanelles-Abella, Keller et al. (2022), Casanelles-Abella, Müller et al. (2022), and Villarroya-Villalba et al. (2021).

**TEXT S2. Additional details on insect sampling and identification.** At each site, trap-nests were installed in trees and in other vertical structures (e.g. lamp post) in the three cases where no trees were available (one in Paris, one in Tartu, and one in Zurich), in other vertical structures. Trap-nests consisted of three pipes. The first two pipes contained 200–300 internodes of the common reed *Phragmites australis* (Cav.) Trin. and 5–10 bamboo internodes (Figure 1b). Reed diameters were between 1 and 10 mm and had a length of 20 cm to cover all requirements of the cavity-nesting bee community. The last pipe was filled only with cardboard tubes of 7.5 mm diameter (Figure 1b), which were specific for large-bodied bees and wasps (WAB Mauerbienenzucht; Konstanz, Germany). Available guides for bees (Amiet et al., 1999, 2002; Falk, 2015), Pompilidae (Wolf, 1972), Sphecidae, Ampulicidae and Crabronidae (Jacobs, 2007), and Eumenidae (Neumeyer, 2019) were used. As in other studies, empty brood cells of eumenid wasps were assumed to belong to the bivoltine *Ancistrocerus nigricornis*, because it was the only species for which offspring of the first generation emerged before trap collection (Fabian et al., 2013; Krewenka et al., 2011).

**TEXT S3.** The local landcover map developed by Alós Ortí et al. (2022) for the same study sites was used to calculate habitat amount. This map has been found to successfully explain bee distribution (Casanelles-Abella, Müller et al., 2022). It distinguishes between bare land, impervious surfaces (buildings and other human-made surfaces), and five types of vegetation (grasslands, shrubs, and coniferous, broadleaf deciduous, and broadleaf evergreen trees). The composition of the different landcover types was calculated within 4, 8, 16 and 32 m buffers from the trap-nest, but due to their correlations only the 32 m buffer was used in this study. In addition, the total tree cover (i.e. combining deciduous and coniferous trees) and woody vegetation cover (i.e. combining shrubs, broadleaved trees and coniferous trees) within a 32 m buffer were calculated.

**TEXT S4.** Land surface temperature (LST) allows us to understand several environmental phenomena, such as thermal pattern of urban landscapes, which can have adverse consequences for wildlife. Cities can create their own microclimate, since the ground is mostly covered by impermeable surfaces, which have a great capacity to absorb solar energy and transform it into heat. This means that throughout the day the impermeable surfaces and therefore the air in the city heats up much more than the vegetated areas. In green areas, a large part of the solar energy is used for evapotranspiration, causing the surface to cool considerably faster. LST is a useful metric for quantifying the thermal stress of bees and wasps since it is influenced by the density of urban fabric, the varied types of urban materials, and the various types of vegetation and its maintenance. LST can be extracted using remotely sensed thermal infrared data.

Several Landsat-8 OLI satellite images were acquired during the most extreme seasons of the year (Table SA) to ensure that LST was not representative of a single day only. These images were obtained from the USGS/Earth Explorer website (https://earthexplorer.usgs.gov/) with a 30 m spatial resolution and, whenever possible, with < 10% cloud cover.

**Text S4 – Table SA.** Remotely sensed thermal infrared data used to determine the land surface temperature (LST) in each of the five cities (Antwerp, Paris, Poznan, Tartu and Zurich).

| City | Sensor | Tile Number | Date | Cloud Coverage |
|---|---|---|---|---|
| Antwerp | Landsat8 - OLI | Path: 198/ Row: 24 | 31/07/2014 | 4.93% |
| Antwerp | Landsat8 - OLI | Path: 198/ Row: 24 | 17/09/2014 | 0.30% |
| Antwerp | Landsat8 - OLI | Path: 198/ Row: 24 | 12/03/2015 | 0.37% |
| Antwerp | Landsat8 - OLI | Path: 198/ Row: 24 | 03/08/2015 | 2.74% |
| Antwerp | Landsat8 - OLI | Path: 198/ Row: 24 | 14/04/2016 | 7.98% |
| Antwerp | Landsat8 - OLI | Path: 198/ Row: 24 | 20/07/2016 | 1.77% |
| Antwerp | Landsat8 - OLI | Path: 198/ Row: 24 | 26/07/2018 | 2.17% |
| Antwerp | Landsat8 - OLI | Path: 198/ Row: 24 | 27/06/2019 | 7.22% |
| Paris | Landsat8 - OLI | Path: 199/ Row: 29 | 16/03/2014 | 8.18% |
| Paris | Landsat8 - OLI | Path: 199/ Row: 26 | 08/09/2014 | 2.36% |
| Paris | Landsat8 - OLI | Path: 199/ Row: 28 | 19/01/2017 | 3.42% |
| Paris | Landsat8 - OLI | Path: 199/ Row: 27 | 23/02/2018 | 0.15% |
| Paris | Landsat8 - OLI | Path: 199/ Row: 26 | 02/08/2018 | 2.82% |
| Paris | Landsat8 - OLI | Path: 199/ Row: 26 | 26/02/2019 | 0.04% |
| Paris | Landsat8 - OLI | Path: 199/ Row: 26 | 04/07/2019 | 0.00% |

| | | | | |
|---|---|---|---|---|
| Paris | Landsat8 - OLI | Path: 199/ Row: 26 | 06/09/2019 | 7.73% |
| Poznan | Landsat8 - OLI | Path: 190/ Row: 24 | 11/08/2015 | 0.03% |
| Poznan | Landsat8 - OLI | Path: 190/ Row: 24 | 14/09/2016 | 0.51% |
| Poznan | Landsat8 - OLI | Path: 190/ Row: 24 | 08/02/2018 | 30.99% |
| Poznan | Landsat8 - OLI | Path: 190/ Row: 24 | 20/09/2018 | 2.30% |
| Poznan | Landsat8 - OLI | Path: 190/ Row: 24 | 27/02/2019 | 8.93% |
| Tartu | Landsat8 - OLI | Path: 187/ Row: 22 | 15/03/2015 | 0.07% |
| Tartu | Landsat8 - OLI | Path: 187/ Row: 19 | 28/12/2015 | 2.35% |
| Tartu | Landsat8 - OLI | Path: 187/ Row: 20 | 26/07/2017 | 11.79% |
| Tartu | Landsat8 - OLI | Path: 186/ Row: 19 | 22/07/2018 | 18.65% |
| Tartu | Landsat8 - OLI | Path: 186/ Row: 19 | 23/08/2018 | 6.05% |
| Tartu | Landsat8 - OLI | Path: 187/ Row: 21 | 17/08/2019 | 29.48% |
| Tartu | Landsat8 - OLI | Path: 187/ Row: 19 | 02/09/2019 | 1.08% |
| Zurich | Landsat8 - OLI | Path: 194/ Row: 24 | 30/08/2015 | 1.29% |
| Zurich | Landsat8 - OLI | Path: 194/ Row: 27 | 18/03/2016 | 1.04% |
| Zurich | Landsat8 - OLI | Path: 194/ Row: 25 | 16/08/2016 | 9.46% |
| Zurich | Landsat8 - OLI | Path: 194/ Row: 26 | 25/08/2016 | 0.51% |
| Zurich | Landsat8 - OLI | Path: 194/ Row: 27 | 08/03/2018 | 8.59% |
| Zurich | Landsat8 - OLI | Path: 195/ Row: 27 | 14/02/2019 | 0.49% |

| | | | | |
|---|---|---|---|---|
| Zurich | Landsat8 - OLI | Path: 195/ Row: 27 | 24/07/2019 | 2.13% |
| Zurich | Landsat8 - OLI | Path: 194/ Row: 27 | 09/08/2019 | 9.42% |
| Zurich | Landsat8 - OLI | Path: 194/ Row: 27 | 18/08/2019 | 0.89% |
| Zurich | Landsat8 - OLI | Path: 195/ Row: 27 | 25/08/2019 | 1.80% |
| Zurich | Landsat8 - OLI | Path: 194/ Row: 27 | 03/09/2019 | 2.95% |

In this work, the radiative transfer equation model was adopted to estimate LST, since it can reach an accuracy of 0.6°C (Sobrino et al., 2004; Yu et al., 2014). It is given by:

$$\mathrm{LST} = \frac{c_2}{\lambda \ln \left\{ \dfrac{c_1}{\lambda^5 \left[ \dfrac{L_{TIR} - Lu - \tau (1 - \varepsilon)\, Ld}{\tau\, \varepsilon} \right]} + 1 \right\}}$$

where LST is the land surface temperature; $c_1$ and $c_2$ are constants; λ is the thermal infrared (TIR) band wavelength; $L_{TIR}$ corresponds to the spectral radiance value at the top-of-atmosphere; Lu and Ld are the upwelling and the downwelling atmospheric radiances, respectively; τ is the atmospheric transmittance; and ε is the land surface emissivity. To simulate atmospheric transmittance and upwelling and downwelling radiances, the spatially and temporally closest atmospheric profile from the MODTRAN model was used. The land surface emissivity was derived based on a normalized difference vegetation index (NDVI) threshold method, using the following equation:

$$\varepsilon = \begin{cases} \varepsilon_w, & \mathrm{NDVI} < 0.0 \\ a + b \cdot \rho_{red}, & 0.0 \geq \mathrm{NDVI} < 0.2 \\ \varepsilon_v \cdot P_v + \varepsilon_s (1 - P_v) + C, & 0.2 \geq \mathrm{NDVI} \leq 0.5 \\ \varepsilon_v + C, & \mathrm{NDVI} > 0.5 \end{cases}$$

where $a$ and b are constants (0.979 and 0.046, respectively); $\rho_{red}$ is the land surface reflectance in the red band; and $\varepsilon_w$, $\varepsilon_v$ and $\varepsilon_s$ are the emissivity of water, vegetation and soil, respectively. $P_v$ refers to the proportion of vegetation and can be calculated using the following equation:

$$P_v = \left[ \frac{\mathrm{NDVI} - \mathrm{NDVI}_{min}}{\mathrm{NDVI}_{max} - \mathrm{NDVI}_{min}} \right]^2$$

where $\mathrm{NDVI}_{min}$ and $\mathrm{NDVI}_{max}$ represent minimum and maximum NDVI, respectively, and can be obtained from the histogram of the NDVI image. C is a term which takes the cavity effect into account, due to the surface roughness, and can be estimated using the following equation:

$$C = (1 - \varepsilon_s) \cdot \varepsilon_v \cdot F \cdot (1 - Pv)$$

where $F$ is the geometrical factor between 0 and 1 (Yu et al., 2014). After processing all indices, the average of each index was calculated for summer and for winter.

ArcMap v.10.5.1 (Esri, Redlands, CA, USA) was used to process Landsat-8 OLI images, including radiometric calibration and atmospheric correction, but also the entire process through the final LST calculation.

In addition to LST, several remote sensing indices were calculated based on Sentinel-2 images. This satellite has a high spatial resolution (10 m) and a 5-day revisit time, therefore offering enormous potential for the analysis of vegetation in urban landscapes (Kopecká et al., 2017). Thus, remote sensing indices derived from Sentinel-2 images can be used as a proxy for the amount of habitat available to urban wild bees. Sentinel-2 images were obtained from the USGS/Earth Explorer website (https://earthexplorer.usgs.gov/), whenever possible with < 10% cloud cover and no clouds under green spaces. For more information about the Sentinel-2 images see Table SB.

**Text S4 – Table SB.** Sentinel-2 data used to determine several remote sensing indices in each of the five cities (Antwerp, Greater Paris, Poznan, Tartu and Zurich).

| City | Sensor | Tile number | Date | Cloud coverage |
|---|---|---|---|---|
| **Antwerp** | Sentinel-2A | T31UES | 12/03/2016 | 3.22% |
| **Antwerp** | Sentinel-2A | T31UES | 20/07/2016 | 0.00% |
| **Antwerp** | Sentinel-2A | T31UES | 27/12/2016 | 0.01% |
| **Antwerp** | Sentinel-2B | T31UES | 25/02/2018 | 0.00% |
| **Antwerp** | Sentinel-2A | T31UES | 30/06/2018 | 0.00% |
| **Antwerp** | Sentinel-2B | T31UES | 15/07/2018 | 0.68% |
| **Antwerp** | Sentinel-2A | T31UES | 18/09/2018 | 6.56% |
| **Antwerp** | Sentinel-2B | T31UES | 21/01/2019 | 6.34% |
| **Antwerp** | Sentinel-2A | T31UES | 15/02/2019 | 0.00% |
| **Antwerp** | Sentinel-2A | T31UES | 25/02/2019 | 0.00% |
| **Antwerp** | Sentinel-2A | T31UES | 25/07/2019 | 3.50% |
| **Antwerp** | Sentinel-2A | T31UES | 24/08/2019 | 2.44% |
| **Paris** | Sentinel-2A | T31UDQ | 16/07/2015 | 0.00% |
| **Paris** | Sentinel-2A | T31UDQ | 27/12/2016 | 0.00% |
| **Paris** | Sentinel-2A | T31UDQ | 26/01/2017 | 0.00% |
| **Paris** | Sentinel-2A | T31UDQ | 15/02/2017 | 0.01% |
| **Paris** | Sentinel-2B | T31UDQ | 25/02/2018 | 0.10% |
| **Paris** | Sentinel-2B | T31UDQ | 25/07/2018 | 0.00% |
| **Paris** | Sentinel-2A | T31UDQ | 19/08/2018 | 5.57% |
| **Paris** | Sentinel-2B | T31UDQ | 21/01/2019 | 0.00% |
| **Paris** | Sentinel-2A | T31UDQ | 15/02/2019 | 0.00% |
| **Paris** | Sentinel-2A | T31UDQ | 25/02/2019 | 0.00% |
| **Poznan** | Sentinel-2A | T33UXU | 10/08/2015 | 0.01% |

| | | | | |
|---|---|---|---|---|
| **Poznan** | Sentinel-2A | T33UXU | 20/08/2015 | 0.02% |
| **Poznan** | Sentinel-2A | T33UXU | 30/07/2017 | 0.00% |
| **Poznan** | Sentinel-2A | T33UXU | 27/12/2017 | 96.11% |
| **Poznan** | Sentinel-2A | T33UXU | 25/02/2018 | 4.45% |
| **Poznan** | Sentinel-2B | T33UXU | 02/03/2018 | 6.95% |
| **Poznan** | Sentinel-2A | T33UXU | 15/07/2018 | 3.41% |
| **Poznan** | Sentinel-2B | T33UXU | 29/08/2018 | 7.50% |
| **Poznan** | Sentinel-2B | T33UXU | 25/02/2019 | 88.55% |
| **Poznan** | Sentinel-2A | T33UXU | 30/06/2019 | 0.00% |
| **Poznan** | Sentinel-2B | T33UXU | 24/08/2019 | 0.00% |
| **Poznan** | Sentinel-2A | T33UXU | 16/01/2020 | 0.00% |
| **Tartu** | Sentinel-2A | T35VME | 04/08/2015 | 0.01% |
| **Tartu** | Sentinel-2A | T35VME | 24/08/2015 | 3.55% |
| **Tartu** | Sentinel-2A | T35VME | 14/02/2017 | 0.06% |
| **Tartu** | Sentinel-2A | T35VME | 06/03/2017 | 0.85% |
| **Tartu** | Sentinel-2A | T35VME | 16/03/2017 | 0.07% |
| **Tartu** | Sentinel-2A | T35VME | 31/07/2017 | 49.73% |
| **Tartu** | Sentinel-2A | T35VME | 30/08/2017 | 0.15% |
| **Tartu** | Sentinel-2A | T35VME | 07/01/2018 | 1.97% |
| **Tartu** | Sentinel-2B | T35VME | 23/08/2018 | 1.52% |
| **Tartu** | Sentinel-2B | T35VME | 19/09/2018 | 0.00% |
| **Tartu** | Sentinel-2B | T35VME | 18/08/2019 | 10.15% |
| **Tartu** | Sentinel-2B | T35VME | 28/08/2019 | 18.82% |
| **Zurich** | Sentinel-2A | T32TMT | 29/08/2015 | 1.52% |
| **Zurich** | Sentinel-2A | T32TMT | 13/08/2016 | 0.00% |
| **Zurich** | Sentinel-2A | T32TMT | 16/02/2017 | 2.36% |
| **Zurich** | Sentinel-2A | T32TMT | 06/07/2017 | 6.54% |
| **Zurich** | Sentinel-2A | T32TMT | 18/08/2017 | 0.44% |
| **Zurich** | Sentinel-2A | T32TMT | 24/07/2018 | 3.75% |
| **Zurich** | Sentinel-2A | T32TMT | 12/09/2018 | 0.05% |
| **Zurich** | Sentinel-2B | T32TMT | 14/02/2019 | 0.07% |
| **Zurich** | Sentinel-2A | T32TMT | 16/02/2019 | 2.79% |
| **Zurich** | Sentinel-2B | T32TMT | 24/02/2019 | 0.78% |
| **Zurich** | Sentinel-2A | T32TMT | 26/02/2019 | 2.44% |
| **Zurich** | Sentinel-2A | T32TMT | 28/08/2019 | 2.07% |

An atmospheric correction was performed using the Sen2Cor plugin (Sen2Cor, v.2.1.2.) from the Sentinel-2 toolbox (SNAP, v.7.0.4). This type of correction is essential because the atmosphere between the satellite and the Earth’s surface reduces the range of possible digital numbers recorded by the sensor.

In addition, it decreases the contrast between adjacent surfaces and changes the brightness of each image pixel. Consequently, spectral indices are, on average, underestimated. This can lead to a weak differentiation of various urban surfaces, becoming a problem in extremely heterogeneous environments.
After completion of the atmospheric correction, the normalized difference vegetation index (NDVI) was estimated with the same software (SNAP, v.7.0.4). This index can be used to characterize existing vegetation and urban infrastructure.
NDVI was developed by Tucker (1979) and is one of the most widely applied indices for monitoring vegetation dynamics. This index results from the following equation:

$$\mathrm{NDVI} = \frac{\rho_{NIR} - \rho_{Red}}{\rho_{NIR} + \rho_{Red}}$$

where $\rho_{NIR}$ and $\rho_{Red}$ are the responses in near-infrared and red bands, respectively. This index represents the photosynthetic capacity, or the energy absorbed by plant canopies, and hence the amount of healthy vegetation. Thus, higher NDVI values indicate a higher density of green vegetation. Specifically, in urban environments, NDVI values > 0.5 correspond to vigorous green areas (Ha and Weng, 2018), while values between 0.2 and 0.5 indicate moisture-stressed vegetation, such as natural meadows. NDVI values near zero and negative values indicate non-vegetated features, such as artificial and barren surfaces, water bodies, snow and clouds.

After all indices were processed, the average of each index for summer and for winter was calculated for each city under study. All indices were computed at different spatial scales (50, 100, 200, 400, 800, 1600 m). ArcMap (v.10.5.1) was used for these calculations.

**Table S1.** Information on the study sites and the samples used. For every site, the area, proximity index and coordinates of the trap-nest position (latitude and longitude) are given.

| City | Site | Area ($m^2$) | Proximity index | Latitude | Longitude |
|---|---|---|---|---|---|
| Antwerp | An011 | 1085854 | 23.57 | 51.16 | 4.36 |
| Antwerp | An016 | 12426 | 931.47 | 51.18 | 4.42 |
| Antwerp | An020 | 20169 | 6.82 | 51.18 | 4.37 |
| Antwerp | An056 | 1054885 | 247.14 | 51.21 | 4.48 |
| Antwerp | An057 | 6704 | 1.52 | 51.21 | 4.39 |
| Antwerp | An062 | 11116 | 3.31 | 51.22 | 4.44 |
| Antwerp | An068 | 93542 | 2.31 | 51.22 | 4.42 |
| Antwerp | An073 | 56928 | 6.78 | 51.22 | 4.39 |
| Antwerp | An082 | 60943 | 4.48 | 51.24 | 4.47 |
| Antwerp | An088 | 14401 | 7.69 | 51.25 | 4.46 |
| Antwerp | An092 | 56167 | 91.92 | 51.26 | 4.45 |
| Antwerp | An102 | 52059 | 3995.62 | 51.29 | 4.43 |
| Paris | Pa282 | 9890 | 3.8 | 48.83 | 2.38 |
| Paris | Pa013 | 126628 | 24.13 | 48.7 | 2.17 |
| Paris | Pa191 | 24993 | 29.42 | 48.8 | 2.3 |
| Paris | Pa245 | 5933064 | 2792.45 | 48.84 | 2.42 |
| Paris | Pa265 | 3553 | 2 | 48.83 | 2.37 |
| Paris | Pa269 | 159611 | 5.39 | 48.82 | 2.34 |
| Paris | Pa295 | 8339 | 5.55 | 48.83 | 2.37 |
| Paris | Pa398 | 169327 | 2.98 | 48.84 | 2.36 |
| Paris | Pa418 | 4630 | 9.83 | 48.84 | 2.29 |
| Paris | Pa492 | 9148 | 45794.28 | 48.85 | 2.26 |
| Paris | Pa535 | 164101 | 49.76 | 48.87 | 2.32 |
| Paris | Pa573 | 4607 | 1.79 | 48.88 | 2.32 |
| Poznan | Po001 | 30443 | 862.43 | 52.31 | 16.98 |
| Poznan | Po037 | 48772 | 11.66 | 52.36 | 16.9 |
| Poznan | Po059 | 8200 | 5.96 | 52.37 | 16.88 |
| Poznan | Po137 | 187103 | 31.09 | 52.39 | 16.93 |
| Poznan | Po179 | 56886 | 3.46 | 52.4 | 16.9 |
| Poznan | Po183 | 10423 | 2136.45 | 52.4 | 16.95 |
| Poznan | Po210 | 13222 | 7.95 | 52.41 | 16.93 |
| Poznan | Po227 | 8406 | 10.5 | 52.41 | 16.87 |
| Poznan | Po267 | 1059825 | 325.97 | 52.43 | 16.95 |
| Poznan | Po348 | 18721 | 18.63 | 52.44 | 16.93 |
| Poznan | Po406 | 5624 | 468.47 | 52.46 | 16.92 |
| Poznan | Po423 | 27974 | 12829.47 | 52.47 | 16.93 |
| Tartu | Ta008 | 6338 | 14.27 | 58.35 | 26.77 |
| Tartu | Ta013 | 2776 | 2.74 | 58.35 | 26.74 |
| Tartu | Ta025 | 6225 | 2.87 | 58.37 | 26.7 |

**TABLE S1.** *Continuation.*

| City | Site | Area ($m^2$) | Proximity index | Latitude | Longitude |
|---|---|---|---|---|---|
| Tartu | Ta033 | 33237 | 5.78 | 58.38 | 26.68 |
| Tartu | Ta040 | 36590 | 314.56 | 58.37 | 26.73 |
| Tartu | Ta047 | 131100 | 57.84 | 58.38 | 26.72 |
| Tartu | Ta057 | 5066 | 5.36 | 58.38 | 26.69 |
| Tartu | Ta064 | 183227 | 14.97 | 58.37 | 26.74 |
| Tartu | Ta102 | 13236 | 22.54 | 58.39 | 26.7 |
| Tartu | Ta104 | 37412 | 5.32 | 58.38 | 26.76 |
| Tartu | Ta110 | 8623 | 7.02 | 58.39 | 26.73 |
| Tartu | Ta125 | 245706 | 26.38 | 58.39 | 26.73 |
| Zurich | Zu006 | 104871 | 104.93 | 47.35 | 8.52 |
| Zurich | Zu007 | 3717 | 7.01 | 47.35 | 8.56 |
| Zurich | Zu015 | 39215 | 167.23 | 47.36 | 8.56 |
| Zurich | Zu018 | 57666 | 56.97 | 47.36 | 8.53 |
| Zurich | Zu033 | 10400 | 28.24 | 47.36 | 8.56 |
| Zurich | Zu039 | 36842 | 10.96 | 47.36 | 8.54 |
| Zurich | Zu057 | 13040 | 6.74 | 47.37 | 8.53 |
| Zurich | Zu062 | 18017 | 6.16 | 47.37 | 8.54 |
| Zurich | Zu067 | 275320 | 14.78 | 47.37 | 8.51 |
| Zurich | Zu080 | 26855 | 8.75 | 47.38 | 8.54 |
| Zurich | Zu082 | 13854 | 17.51 | 47.38 | 8.49 |
| Zurich | Zu087 | 22711 | 4.87 | 47.39 | 8.52 |
| Zurich | Zu094 | 96182 | 974.64 | 47.39 | 8.47 |
| Zurich | Zu105 | 9576 | 67.97 | 47.4 | 8.5 |
| Zurich | Zu113 | 46486 | 34334.06 | 47.4 | 8.52 |
| Zurich | Zu119 | 107938 | 108059.38 | 47.4 | 8.54 |
| Zurich | Zu126 | 11737 | 15.67 | 47.4 | 8.5 |
| Zurich | Zu133 | 3511 | 13.91 | 47.41 | 8.54 |
| Zurich | Zu155 | 4346 | 6.51 | 47.41 | 8.55 |
| Zurich | Zu158 | 5929 | 7.75 | 47.41 | 8.53 |
| Zurich | Zu173 | 5600 | 25.03 | 47.42 | 8.51 |
| Zurich | Zu179 | 103083 | 2778.23 | 47.42 | 8.53 |
| Zurich | Zu901 | 15459 | 32.08 | 47.41 | 8.48 |
| Zurich | Zu903 | 58342 | 332.17 | 47.41 | 8.51 |
| Zurich | Zu904 | 8253 | 5.04 | 47.39 | 8.52 |
| Zurich | Zu905 | 10987 | 7.02 | 47.41 | 8.56 |
| Zurich | Zu906 | 10629 | 9.1 | 47.4 | 8.59 |
| Zurich | Zu907 | 22894 | 25.21 | 47.4 | 8.49 |
| Zurich | Zu908 | 102401 | 262.43 | 47.35 | 8.58 |
| Zurich | Zu910 | 53898 | 14.5 | 47.34 | 8.53 |
| Zurich | Zu911 | 3219 | 18.09 | 47.43 | 8.5 |
| Zurich | Zu912 | 26567 | 8.86 | 47.35 | 8.55 |

**TABLE S2.** List of cavity-nesting bee and wasp species collected in the 80 study sites within 5 cities. For every species, the total number of brood cells is provided.

| | | | |
|---|---|---|---|
| **Bees** | **Colletidae** | | |
| | | *Hylaeus communis* (Nylander, 1852) | 1007 |
| | | *Hylaeus confusus* (Nylander, 1852) | 35 |
| | | *Hylaeus incongruus* (Förster 1871) | 5 |
| | | *Hylaeus pectoralis* (Förster, 1871) | 12 |
| | | *Hylaeus punctulatissimus* (Smith 1842) | 4 |
| | | *Hylaeus* sp. | 748 |
| | **Megachillidae** | | |
| | | *Anthidium manicatum* (Linnaeus, 1758) | 13 |
| | | *Chelostoma campanularum* (Lepeletier, 1841) | 21 |
| | | *Chelostoma florisomne* (Linnaeus, 1758) | 1728 |
| | | *Chelostoma rapunculi* (Lepeletier, 1841) | 24 |
| | | *Chelostoma* sp. | 36 |
| | | *Heriades truncorum* (Linnaeus, 1758) | 180 |
| | | *Megachile centuncularis* (Linnaeus, 1758) | 98 |
| | | *Megachile* sp. | 71 |
| | | *Osmia bicornis* (Linnaeus, 1758) | 3173 |
| | | *Osmia brevicornis* (Fabricius, 1798) | 8 |
| | | *Osmia caerulescens* (Linnaeus, 1758) | 93 |
| | | *Osmia cornuta* | 1230 |
| | | *Osmia leaiana* (Kirby, 1802) | 26 |
| | | *Osmia* sp. | 401 |
| | | Undetermined | 4 |
| **Wasps** | **Ampulicidae** | | |
| | | *Ampulex fasciata* (Jurine, 1807) | 31 |
| | | *Dolichurus corniculatus* (Spinola, 1808) | 1 |
| | **Crabronidae** | | |
| | | *Nitela borealis* (Valkeila, 1974) | 182 |
| | | *Nitela spinolae* (Latreille, 1809) | 38 |
| | | *Nitela* sp. | 327 |
| | | *Passaloecus borealis* (Dahlbom, 1844) | 113 |
| | | *Passaloecus brevilabris* (Wolf, 1958) | 19 |
| | | *Passaloecus corniger* (Shuckard, 1837) | 190 |
| | | *Passaloecus eremita* (Kohl, 1893) | 778 |
| | | *Passaloecus gracilis* (Kohl, 1893) | 369 |
| | | *Passaloecus insignis* (Vander Linden, 1829) | 517 |
| | | *Passaloecus monilicornis* (Dahlbom, 1842) | 28 |
| | | *Passaloecus pictus* (Ribaut, 1952) | 1 |
| | | *Passaloecus* sp. | 1118 |
| | | *Pemphredon lugens* | 52 |
| | | *Pemphredon lugubris* | 3 |
| | | *Pemphredon mortifer* | 5 |
| | | *Pemphredon* sp. | 11 |
| | | *Pison* sp. | 6 |
| | | *Psenulus fuscipennis* (Vikberg, 1986) | 135 |
| | | *Psenulus pallipes* (Panzer, 1798) | 33 |
| | | *Psenulus* sp. | 28 |
| | | *Rhopalum clavipes* (Linnaeus, 1758) | 7 |
| | | *Spilomena troglodytes* (Vander Linden, 1829) | 244 |
| | | *Spilomena* sp. | 37 |
| | | *Trypoxylon attenuatum* (Smith, 1851) | 1 |
| | | *Trypoxylon clavicerum* (Lepeletier & Serville, 1828) | 101 |
| | | *Trypoxylon figulus* (Linneaus, 1758) | 74 |
| | | *Trypoxylon fronticorne* (Gussakovskij, 1936) | 4 |
| | | *Trypoxylon kolazyi* (Kohl, 1893) | 64 |
| | | *Trypoxylon minus* (de Beaumont, 1945) | 64 |
| | | *Trypoxylon* sp. | 341 |
| | | Undetermined | 66 |
| | **Pompilidae** | | |
| | | *Agenioideus cinctellus* (Spinola, 1807) | 42 |
| | | *Agenioideus* sp. | 22 |
| | | *Anoplius concinnus* (Dahlbom, 1845) | 1 |
| | | *Auplopus carbonarius* (Scopoli, 1763) | 76 |
| | | *Auplopus* sp. | 2 |
| | | *Dipogon bifasciatus* (Geoffroy, 1785) | 4 |
| | | *Dipogon hircanum* (Fabricius, 1798) | 3 |
| | | *Dipogon* sp. | 149 |
| | | *Dipogon subintermedius* (Magretti, 1886) | 361 |
| | | *Dipogon variegatus* (Linnaeus, 1758) | 47 |
| | | Pompilidae sp. | 55 |
| | | undetermined | |
| | **Sphecidae** | | |
| | | *Isodontia mexicana* (Saussure 1867) | 10 |
| | **Vespidae** | | |
| | | *Allodynerus rossii* (Lepeletier, 1841) | 3 |
| | | *Ancistrocerus antilope* (Panzer, 1798) | 69 |
| | | *Ancistrocerus claripennis* (Giordani Soika, 1970) | 13 |
| | | *Ancistrocerus gazella* (Panzer, 1798) | 13 |
| | | *Ancistrocerus nigricornis* (Curtis, 1826) | 1122 |
| | | *Ancistrocerus* sp. | 11 |
| | | *Ancistrocerus trifasciatus* (Müller, 1776) | 191 |
| | | *Microdynerus timidus* (Saussure, 1856) | 12 |
| | | *Symmorphus angustatus* (Zetterstedt, 1838) | 53 |
| | | *Symmorphus bifasciatus* (Linnaeus, 1761) | 93 |
| | | *Symmorphus connexus* (Curtis, 1826) | 76 |
| | | *Symmorphus crassicornis* (Panzer, 1798) | 132 |
| | | *Symmorphus* sp. | 16 |
| | | Undetermined | 103 |

**TABLE S3.** Natural enemy species collected in the 80 study sites within 5 cities. Natural enemies are sorted according to their order. For every natural enemy, the host species list and the host order (O) are provided with the number of parasitized brood cells (N). B = bees; O = undetermined; W = wasps.

| Enemy order and species | Host species | O | N | Enemy order and species | Host species | O | N |
|---|---|---|---|---|---|---|---|
| **Acari** | | | | ***Trichodes apiarius* (Linnaeus, 1758)** | | | |
| ***Chaetodactylus* sp. *** | | | | | *Osmia sp.* | B | 1 |
| | *Osmia cornuta* | B | 7 | ***Trogoderma glabrum* (Herbst, 1783)** | | | |
| **Coleoptera** | | | | | *Osmia bicornis* | B | 2 |
| ***Anthrenus verbasci* (Linnaeus, 1767)** | | | | **Diptera** | | | |
| | *Chelostoma florisomne* | B | 1 | ***Anthrax anthrax* (Schrank, 1781)** | | | |
| | *Isodontia mexicana* | W | 1 | | *Osmia bicornis* | B | 1 |
| | *Nitela sp.* | W | 2 | | *Osmia sp.* | B | 1 |
| | *Osmia caerulescens* | B | 5 | ***Cacoxenus indagator* (Loew, 1858)** | | | |
| | *unknown wasp* | W | 1 | | *Chelostoma florisomne* | B | 1 |
| **Dermestidae sp.** | | | | | *Osmia bicornis* | B | 242 |
| | *Dipogon sp.* | W | 1 | | *Osmia caerulescens* | B | 1 |
| | *Eumeninae sp.* | W | 13 | | *Osmia cornuta* | B | 65 |
| | *Hylaeus sp.* | B | 1 | | *Osmia sp.* | B | 70 |
| | *Isodontia mexicana* | W | 2 | **Tachinidae sp. *** | | | |
| | *Nitela sp.* | W | 3 | | *Ancistrocerus trifasciatus* | W | 5 |
| | *Osmia bicornis* | B | 1 | | *Eumeninae sp.* | W | 33 |
| | *Osmia sp.* | B | 1 | | *Megachile sp.* | B | 1 |
| | *Passaloecus gracilis* | W | 2 | | *Osmia sp.* | B | 3 |
| | *Passaloecus sp.* | W | 1 | | *Pompilidae sp.* | W | 1 |
| | *Pompilidae sp.* | W | 3 | | *Symmorphus sp.* | W | 1 |
| ***Megatoma undata* (Linnaeus, 1752)** | | | | | *Trypoxylon sp.* | W | 1 |
| | *Anthidium manicatum* | B | 2 | | *unknown wasp* | W | 1 |
| | *Crabronidae sp.* | W | 1 | **Braconidae sp. *** | | | |
| | *Dipogon sp.* | W | 2 | | *Dipogon subintermedius* | W | 1 |
| | *Eumeninae sp.* | W | 24 | | *Eumeninae sp.* | W | 7 |
| | *Hylaeus sp.* | B | 5 | | *Trypoxylon sp.* | W | 2 |
| | *Osmia bicornis* | B | 8 | **Chrysidae sp. *** | | | |
| | *Osmia cornuta* | B | 9 | | *Eumeninae sp.* | W | 1 |
| | *Osmia leaiana* | B | 3 | | *Passaloecus eremita* | W | 2 |
| | *Osmia sp.* | B | 35 | | *Passaloecus insignis* | W | 1 |
| | *Passaloecus sp.* | W | 7 | | *Passaloecus sp.* | W | 2 |
| | *Pompilidae sp.* | W | 2 | | *Pemphredon lugens* | W | 2 |
| | *Trypoxylon sp.* | W | 1 | | *Symmorphus bifasciatus* | W | 1 |
| | *unknown wasp* | W | 5 | | *Trypoxylon sp.* | W | 2 |
| ***Ptinus sexpunctatus* (Panzer, 1789)** | | | | ***Chrysis* cf. *angustula* (Schenck, 1856)** | | | |
| | *Chelostoma florisomne* | B | 2 | | *Symmorphus bifasciatus* | W | 3 |
| | *Chelostoma sp.* | B | 1 | ***Chrysis* cf. *Ruddii* (Shuckard, 1837)** | | | |
| | *Eumeninae sp.* | W | 1 | | *Symmorphus bifasciatus* | W | 4 |
| | *Hylaeus sp.* | B | 3 | | *Symmorphus crassicornis* | W | 3 |
| | *Osmia bicornis* | B | 1 | ***Chrysis fulgida* (Linnaeus, 1761)** | | | |
| | *Osmia sp.* | B | 4 | | *Eumeninae sp.* | W | 3 |
| | *Pemphredon lugens* | W | 1 | | *Symmorphus angustatus* | W | 13 |

**TABLE S3.** *Continuation.*

| Enemy order and species | Host species | O | N | Enemy order and species | Host species | O | N |
|---|---|---|---|---|---|---|---|
| ***Chrysis ignita*** **(Linnaeus, 1758)** | | | | ***Gasteruption*** **sp.** | | | |
| | *Ancistrocerus trifasciatus* | W | 3 | | *Hylaeus sp.* | B | 3 |
| | *Symmorphus bifasciatus* | W | 2 | **Hymenoptera sp. *** | | | |
| ***Chrysis*** **sp.** | | | | | *Auplopus carbonarius* | W | 8 |
| | *Eumeninae sp.* | W | 1 | | *Dipogon sp.* | W | 1 |
| ***Coelioxys inermis*** **(Kirby 1802)** | | | | | *Eumeninae sp.* | W | 3 |
| | *Megachile centuncularis* | B | 1 | | *Ancistrocerus claripennis* | W | 3 |
| ***Coelopencyrtus*** **sp.** | | | | | *Ancistrocerus trifasciatus* | W | 1 |
| | *Hylaeus communis* | B | 5 | | *Nitela sp.* | W | 1 |
| **Dermestidae sp.** | | | | | *Osmia bicornis* | B | 6 |
| | *Osmia sp.* | B | 1 | **Hymenopter** | *Osmia brevicornis* | B | 1 |
| ***Ephialtes manifestator*** **(Linnaeus, 1758)** | | | 1 | | *Osmia sp.* | B | 3 |
| | *Spilomena sp.* | W | 7 | | *Passaloecus gracilis* | W | 1 |
| | *Spilomena troglodytes* | W | | | *Passaloecus sp.* | W | 1 |
| ***Ephialtes*** **sp. *** | | | 1 | | *Trypoxylon sp.* | W | 2 |
| | *Passaloecus insignis* | W | | ***Ichneumon*** **sp. *** | | | |
| **Eulophidae sp. *** | | | 1 | | *Hylaeus sp.* | B | 1 |
| | *Crabronidae sp.* | W | | | *Megachillidae sp.* | B | 1 |
| ***Eurytoma nodularis*** **(Boheman, 1836)** | | | | | *Passaloecus borealis* | W | 1 |
| | *Auplopus carbonarius* | W | 2 | | *Passaloecus eremita* | W | 5 |
| | *Dipogon sp.* | W | 2 | | *Passaloecus insignis* | W | 1 |
| | *Hylaeus sp.* | B | 2 | | *Passaloecus sp.* | W | 7 |
| | *Nitela sp.* | W | 1 | | *Pompilidae sp.* | W | 2 |
| | *Osmia sp.* | B | 1 | | *Psenulus fuscipennis* | W | 2 |
| | *Pison sp.* | W | 1 | | *Symmorphus crassicornis* | W | 1 |
| | *Symmorphus sp.* | W | 1 | | *Trypoxylon figulus* | W | 1 |
| | *Trypoxylon sp.* | W | 1 | | *Trypoxylon minus* | W | 2 |
| **Eurytomidae sp. *** | | | | | *Trypoxylon sp.* | W | 1 |
| | *Ampulex fasciata* | W | 5 | | *unknown wasp* | W | 1 |
| | *Dipogon sp.* | W | 1 | ***Lochetica*** **sp. *** | | | |
| | *Hylaeus sp.* | B | 3 | | *Passaloecus insignis* | W | 1 |
| | *Nitela borealis* | W | 3 | ***Melittobia acasta*** **(Walker, 1839)** | | | |
| | *Nitela sp.* | W | 11 | | *Agenioideus cinctellus* | W | 3 |
| | *Trypoxylon sp.* | W | 1 | | *Ampulex fasciata* | W | 7 |
| ***Gasteruption assectator*** **agg.** | | | | | *Auplopus carbonarius* | W | 3 |
| | *Eumeninae sp.* | W | 1 | | *Chelostoma florisomne* | B | 2 |
| | *Hylaeus communis* | B | 9 | | *Crabronidae sp.* | W | 1 |
| | *Hylaeus sp.* | B | 41 | | *Dipogon sp.* | W | 25 |
| | *Trypoxylon sp.* | W | 1 | | *Eumeninae sp.* | W | 11 |
| ***Gasteruption jaculator*** **(Linnaeus, 1758)** | | | | | *Heriades truncorum* | B | 1 |
| | *Hylaeus confusus* | B | 1 | | *Hylaeus sp.* | B | 5 |
| | *Hylaeus sp.* | B | 8 | | *Isodontia mexicana* | W | 3 |
| | | | | | *Megachile sp.* | B | 28 |

**TABLE S3.** *Continuation.*

| Enemy order and species | Host species | O | N |
|---|---|---|---|
| | *Nitela sp.* | W | 69 |
| | *Osmia cornuta* | B | 7 |
| | *Passaloecus gracilis* | W | 8 |
| | *Passaloecus sp.* | W | 94 |
| | *Pompilidae sp.* | W | 1 |
| | *Spilomena sp.* | W | 1 |
| | *Trypoxylon minus* | W | 5 |
| | *Trypoxylon sp.* | W | 66 |
| ***Monodontomerus obscurus (Westwood, 1833)*** | | | |
| | *Auplopus carbonarius* | W | 4 |
| | *Osmia bicornis* | B | 12 |
| | *Osmia cornuta* | B | 12 |
| | *Osmia sp.* | B | 16 |
| | *Trypoxylon sp.* | W | 1 |
| ***Monodontomerus* sp. *** | | | |
| | *Osmia bicornis* | B | 18 |
| | *Osmia cornuta* | B | 9 |
| | *Osmia sp.* | B | 1 |
| | *Passaloecus* sp. | W | 1 |
| | *unknown wasp* | W | 2 |
| ***Monosapyga clavicornis* (Linnaeus, 1758)** | | | |
| | *Chelostoma florisomne* | B | 66 |
| ***Neorhacodes enslini (Ruschka, 1922)*** | | | |
| | *Spilomena troglodytes* | W | 10 |
| ***Omalus aeneus*** | | | |
| | *Crabronidae sp.* | W | 2 |
| | *Passaloecus eremita* | W | 11 |
| | *Passaloecus gracilis* | W | 8 |
| | *Passaloecus insignis* | W | 3 |
| | *Passaloecus sp.* | W | 21 |
| | *unknown* | O | 1 |
| ***Omalus biaccinctus*** | | | |
| | *Passaloecus borealis* | W | 1 |
| | *Passaloecus sp.* | W | 3 |
| ***Omalus puncticollis*** | | | |
| | *Passaloecus eremita* | W | 8 |
| | *Pompilidae sp.* | W | 1 |
| ***Omalus pusillus*** | | | |
| | *Passaloecus eremita* | W | 1 |
| ***Perithous scurra*** | | | |
| | *Psenulus pallipes* | W | 1 |
| ***Poemenia* sp. *** | | | |
| | *Dipogon subintermedius* | W | 1 |
| | *Passaloecus gracilis* | W | 1 |
| | *Passaloecus monilicornis* | W | 3 |
| | *Psenulus fuscipennis* | W | 7 |
| | *Psenulus pallipes* | W | 2 |
| | *Trypoxylon sp.* | W | 1 |
| ***Pseudomalus auratus*** | | | |
| | *Ancistrocerus nigricornis* | W | 1 |
| | *Crabronidae sp.* | W | 1 |
| | *Hylaeus communis* | B | 1 |
| | *Nitela sp.* | W | 1 |
| | *Passaloecus corniger* | W | 8 |
| | *Passaloecus eremita* | W | 28 |
| | *Passaloecus gracilis* | W | 3 |
| | *Passaloecus insignis* | W | 2 |
| | *Passaloecus monilicornis* | | 1 |
| | *Passaloecus sp.* | W | 19 |
| | *Psenulus fuscipennis* | W | 1 |
| | *Psenulus sp.* | W | 1 |
| ***Pseudomalus pusillus*** | | | |
| | *Eumeninae sp.* | W | 1 |
| | *Passaloecus insignis* | W | 1 |
| ***Pseudomalus* sp. *** | | | |
| | *Passaloecus sp.* | W | 1 |
| ***Pseudomalus triangulifer*** | | | |
| | *Nitela borealis* | W | 1 |
| | *Passaloecus eremita* | W | 25 |
| | *Passaloecus sp.* | W | 1 |
| | *Pemphredon lugens* | W | 1 |
| ***Pseudomalus violaceus*** | | | |
| | *Pemphredon lugens* | W | 2 |
| ***Pteromalidae* sp.** | | | |
| | *Dipogon subintermedius* | W | 1 |
| ***Sapyga quinquepunctata*** | | | |
| | *Osmia caerulescens* | B | 4 |
| ***Trichrysis cyanea*** | | | |
| | *Dipogon subintermedius* | W | 1 |
| | *Psenulus sp.* | W | 2 |
| | *Symmorphus crassicornis* | W | 1 |
| | *Trypoxylon figulus* | W | 2 |
| | *Trypoxylon sp.* | W | 2 |

**TABLE S4.** Summary of the proxies used for ecological drivers (i.e. habitat amount, resource availability, temperature, top-down control and bottom-up control). The relationship between the proxy and the actual driver is positive (green) or negative (red).

| Type | Proxy | Driver | | |
|---|---|---|---|---|
| | | **Amount of available habitat** | **Temperature** | **Biotic interactions** |
| Spatial analysis | Proximity index at 500 m | | | |
| Spatial analysis | Site size | | | |
| Remote sensing | NDVI at 100 and 800 m | | | |
| Mapping | Grassland cover at 32 m | | | |
| Mapping | Tree cover 32 m | | | |
| Mapping | Woody vegetation cover at 32 m | | | |
| Mapping | Impervious surface cover at 32 m | | | |
| Field | Plant species richness at 100 m | | | |
| Field | Plant Shannon diversity at 100 m | | | |
| Remote sensing | LST at 800 | | | |
| Sensor | Local temperature | | | |
| Trap-nest | Parasitism rate | | | |
| Trap-nest | Number of nests | | | |

**TABLE S5.** Summary of the sampled cavity-nesting bees and wasps in the five studied cities. For each city and taxonomic group, statistics relating to the community structure, community composition, performance and life-history traits are provided.

| | Antwerp | | Paris | | Poznan | | Tartu | | Zurich | |
|---|---|---|---|---|---|---|---|---|---|---|
| **Response** | **Bees** | **Wasps** | **Bees** | **Wasps** | **Bees** | **Wasps** | **Bees** | **Wasps** | **Bees** | **Wasps** |
| Species richness | 6 | 22 | 12 | 22 | 5 | 18 | 5 | 17 | 11 | 34 |
| Enemy richness | 7 | 7 | 8 | 12 | 8 | 16 | 4 | 15 | 17 | 16 |
| Total N. nests | 267 | 306 | 588 | 463 | 210 | 231 | 101 | 189 | 832 | 1205 |
| Total N. cells | 1227 | 1016 | 2508 | 1202 | 1098 | 876 | 537 | 897 | 3547 | 3709 |
| Total parasitized | 135 | 158 | 180 | 89 | 108 | 103 | 32 | 45 | 289 | 361 |
| Total hatched | 760 | 341 | 1517 | 375 | 519 | 319 | 294 | 534 | 2200 | 1171 |
| Total died | 467 | 675 | 991 | 827 | 579 | 557 | 243 | 363 | 1347 | 2538 |
| Total females | 425 | 194 | 510 | 217 | 263 | 211 | 240 | 36 | 1251 | 710 |
| Total males | 360 | 135 | 1011 | 160 | 298 | 137 | 66 | 206 | 1076 | 509 |
| Average N. cells per nest (min - max) | 4.60 (1, 12) | 3.29 (1, 20) | 4.25 (1, 13) | 2.60 (1, 19) | 5.23 (1, 19) | 3.78 (1, 14) | 5.32 (1, 19) | 4.70 (1, 20) | 4.25 (1, 21) | 3.06 (1, 23) |
| Average N. females per nest (min - max) | 1.59 (0, 10) | 0.63 (0, 10) | 0.86 (0, 8) | 0.47 (0, 7) | 1.25 (0, 16) | 0.91 (0, 10) | 2.38 (0, 11) | 1.76 (0, 19) | 1.50 (0, 10) | 0.57 (0, 7) |
| Average N. males per nest (min - max) | 1.35 (0, 6) | 0.44 (0, 17) | 1.71 (0, 10) | 0.35 (0, 8) | 1.42 (0, 9) | 0.59 (0, 7) | 0.65 (0, 6) | 1.08 (0, 11) | 1.29 (0, 9) | 0.42 (0, 6) |
| Average N. dead cells per nest (min - max) | 1.72 (0, 9) | 1.51 (0,11) | 1.67 (0, 11) | 1.6 (0, 18) | 2.71 (0, 18) | 1.47 (0, 10) | 2.41 (0, 17) | 1.72 (0, 10) | 1.60 (0, 15) | 1.70 (0, 18) |
| Average N. Cells with enemies per nest (min - max) | 0.51 (0, 9) | 0.51 0, 10) | 0.31 (0, 10) | 0.19 (0, 6) | 0.51 (0, 8) | 044 (0, 8) | 0.32 (0, 6) | 0.24 (0, 4) | 0.34 (0, 7) | 0.30 (0, 16) |
| Most common genus | *Chelostoma* spp. | *Trypoxylon* spp. | *Osmia* spp. | *Nitela* spp. | *Osmia* spp. | *Symmorphus* spp. | *Hylaeus* spp. | *Passaloecus* spp. | *Osmia* spp. | *Passaloecus* spp. |
| Most common species | *Osmia cornuta* | *Trypoxylon clavicerum* | *Osmia bicornis* | *Nitela borealis* | *Osmia bicornis* | *Symmorphus crassicornis* | *Hylaeus communis* | *Spilomena troglodytes* | *Chelostoma florisomne* | *Passaloecus eremita* |

**TABLE S6.** Output of the multimodal inference. For every response within a group, the estimates, standard errors (se), z-values (Statistic) and p-values are provided. Significance is set at $p < 0.05$.

| Group | Response | Predictors | Estimate ± se | Statistic | p-value | |
|---|---|---|---|---|---|---|
| Bees | Species richness | Intercept | 0.710 ± 0.097 | 7.211 | <0.001 | *** |
| | | Local temperature | 0.024 ± 0.066 | 0.358 | 0.721 | |
| | | LST 800 | -0.001 ± 0.025 | 0.025 | 0.958 | |
| | | Park area | -0.001 ± 0.023 | 0.033 | 0.974 | |
| | | Proximity index | -0.002 ± 0.022 | 0.099 | 0.921 | |
| | | NDVI 100 | 0.286 ± 0.094 | 2.985 | 0.002 | ** |
| | | NDVI 800 | -0.003 ± 0.025 | 0.101 | 0.919 | |
| | | Grasslands 32 | -0.001 ± 0.019 | 0.067 | 0.947 | |
| | | Woody 32 | 0.019 ± 0.056 | 0.336 | 0.737 | |
| | | Impervious 32 | -0.002 ± 0.022 | 0.075 | 0.940 | |
| | | Plant S | 0.045 ± 0.079 | 0.572 | 0.567 | |
| | | Plant H' | 0.176 ± 0.132 | 1.328 | 0.184 | |
| | | Parasitism rate | 0.087 ± 0.367 | 0.236 | 0.813 | |
| | | N. parasites | 0.001 ± 0.003 | 0.324 | 0.746 | |
| | Abundance hatched | Intercept | 3.958 ± 0.268 | 14.519 | <0.001 | *** |
| | | Local temperature | 0.182 ± 0.281 | 0.645 | 0.519 | |
| | | LST 800 | -0.065 ± 0.017 | 0.394 | 0.694 | |
| | | Park area | 0.003 ± 0.025 | 0.106 | 0.915 | |
| | | Proximity index | 0.009 ± 0.046 | 0.187 | 0.851 | |
| | | NDVI 100 | 0.563 ± 0.263 | 2.123 | 0.034 | * |
| | | NDVI 800 | 0.338 ± 0.263 | 1.277 | 0.202 | |
| | | Plant S | 0.045 ± 0.126 | 0.358 | 0.720 | |
| | | Plant H' | 0.301 ± 0.278 | 1.078 | 0.281 | |
| Wasps | Species richness | Intercept | 1.257 ± 0.116 | 10.651 | <0.001 | *** |
| | | Local temperature | 0.008 ± 0.038 | 0.209 | 0.835 | |
| | | LST 800 | 0.000 ± 0.011 | 0.029 | 0.977 | |
| | | Park area | 0.000 ± 0.007 | 0.054 | 0.957 | |
| | | Proximity index | -0.053 ± 0.110 | 0.48 | 0.631 | |
| | | NDVI 100 | 0.324 ± 0.068 | 4.669 | <0.001 | *** |
| | | NDVI 800 | -0.002 ± 0.019 | 0.126 | 0.900 | |
| | | Grasslands 32 | -0.052 ± 0.080 | 0.649 | 0.516 | |
| | | Woody 32 | 0.017 ± 0.049 | 0.345 | 0.730 | |
| | | Impervious 32 | 0.001 ± 0.011 | 0.065 | 0.948 | |
| | | Plant S | 0.000 ± 0.008 | 0.046 | 0.964 | |
| | | Plant H' | 0.001 ± 0.011 | 0.054 | 0.957 | |
| | | Parasitism rate | 0.013 ± 0.004 | 2.95 | 0.003 | ** |
| | | N. parasites | -0.037 ± 0.205 | 0.18 | 0.857 | |
| | Abundance hatched | Intercept | 3.440 ± 0.146 | 23.183 | <0.001 | *** |
| | | Local temperature | 0.001 ± 0.044 | 0.032 | 0.975 | |
| | | LST 800 | 0.000 ± 0.047 | 0.01 | 0.992 | |
| | | Park area | 0.002 ± 0.047 | 0.043 | 0.966 | |
| | | Proximity index | -0.128 ± 0.199 | 0.638 | 0.523 | |
| | | NDVI 100 | 0.577 ± 0.156 | 3.643 | <0.001 | *** |
| | | NDVI 800 | -0.006 ± 0.063 | 0.092 | 0.927 | |
| | | Grasslands 32 | 0.003 ± 0.046 | 0.057 | 0.954 | |
| | | Woody 32 | 0.014 ± 0.065 | 0.213 | 0.831 | |
| | | Impervious 32 | -0.006 ± 0.050 | 0.125 | 0.901 | |
| | | Plant S | -0.002 ± 0.047 | 0.042 | 0.967 | |
| | | Plant H' | 0.006 ± 0.053 | 0.107 | 0.914 | |
| | | Parasitism rate | -0.210 ± 0.659 | 0.315 | 0.753 | |

**TABLE S6.** *Continuation*

| Group | Response | Predictors | Estimate ± se | | | Statistic | p-value | |
|---|---|---|---|---|---|---|---|---|
| Bees | Survival | Intercept | 0.066 | ± | 0.159 | 0.414 | 0.679 | |
| | | Local temperature | 0.026 | ± | 0.094 | 0.272 | 0.786 | |
| | | LST 50 | -0.020 | ± | 0.097 | 0.207 | 0.836 | |
| | | Park area | 0.009 | ± | 0.048 | 0.179 | 0.858 | |
| | | Proximity index | 0.062 | ± | 0.127 | 0.486 | 0.627 | |
| | | NDVI 100 | 0.217 | ± | 0.293 | 0.742 | 0.458 | |
| | | NDVI 800 | 0.630 | ± | 0.278 | 2.265 | 0.024 | * |
| | | Grasslands 32 | 0.004 | ± | 0.054 | 0.082 | 0.935 | |
| | | Woody 32 | -0.024 | ± | 0.086 | 0.28 | 0.780 | |
| | | Impervious 32 | 0.015 | ± | 0.074 | 0.204 | 0.838 | |
| | | Plant S | 0.005 | ± | 0.082 | 0.061 | 0.952 | |
| | | Plant H' | 0.081 | ± | 0.157 | 0.518 | 0.605 | |
| | Parasitism | Intercept | -8.288 | ± | 0.288 | 28.759 | <0.001 | *** |
| | | Local temperature | 0.011 | ± | 0.195 | 0.055 | 0.956 | |
| | | LST 800 | 0.052 | ± | 0.227 | 0.916 | 0.359 | |
| | | Park area | 0.006 | ± | 0.068 | 0.418 | 0.676 | |
| | | Proximity index | 0.008 | ± | 0.129 | 0.336 | 0.737 | |
| | | NDVI 100 | -0.019 | ± | 0.201 | 0.483 | 0.629 | |
| | | NDVI 800 | 0.050 | ± | 0.189 | 0.264 | 0.792 | |
| | | Grasslands 32 | 0.014 | ± | 0.223 | 0.332 | 0.740 | |
| | | Woody 32 | 0.014 | ± | 0.226 | 0.323 | 0.746 | |
| | | Impervious 32 | -0.031 | ± | 0.211 | 0.668 | 0.504 | |
| | | Plant S | -0.003 | ± | 0.112 | 0.133 | 0.894 | |
| | | Plant H' | -0.010 | ± | 0.155 | 0.332 | 0.740 | |
| | Females | Intercept | 0.305 | ± | 0.124 | 2.466 | 0.014 | * |
| | | Local temperature | -0.703 | ± | 0.141 | 4.997 | <0.001 | *** |
| | | LST 50 | -0.004 | ± | 0.049 | 0.074 | 0.941 | |
| | | Park area | -0.026 | ± | 0.065 | 0.403 | 0.687 | |
| | | Proximity index | 0.000 | ± | 0.029 | 0.017 | 0.987 | |
| | | NDVI 100 | -0.025 | ± | 0.088 | 0.280 | 0.780 | |
| | | NDVI 800 | -0.147 | ± | 0.171 | 0.858 | 0.391 | |
| | | Grasslands 32 | -0.016 | ± | 0.062 | 0.254 | 0.799 | |
| | | Woody 32 | 0.007 | ± | 0.047 | 0.148 | 0.882 | |
| | | Impervious 32 | 0.019 | ± | 0.067 | 0.283 | 0.777 | |
| | | Plant S | 0.003 | ± | 0.042 | 0.065 | 0.948 | |
| | | Plant H' | -0.017 | ± | 0.069 | 0.253 | 0.800 | |
| | Number of cells | Intercept | 1.497 | ± | 0.048 | 31.191 | <0.001 | *** |
| | | Local temperature | -0.051 | ± | 0.040 | 1.269 | 0.204 | |
| | | LST 50 | -0.023 | ± | 0.040 | 0.585 | 0.558 | |
| | | Park area | 0.002 | ± | 0.019 | 0.081 | 0.935 | |
| | | Proximity index | 0.001 | ± | 0.021 | 0.035 | 0.972 | |
| | | NDVI 100 | 0.001 | ± | 0.037 | 0.014 | 0.989 | |
| | | NDVI 800 | 0.107 | ± | 0.030 | 3.528 | <0.001 | *** |
| | | Grasslands 32 | 0.027 | ± | 0.028 | 0.958 | 0.338 | |
| | | Woody 32 | -0.020 | ± | 0.031 | 0.642 | 0.521 | |
| | | Impervious 32 | -0.042 | ± | 0.029 | 1.440 | 0.150 | |
| | | Plant S | 0.002 | ± | 0.024 | 0.063 | 0.950 | |
| | | Plant H' | 0.011 | ± | 0.043 | 0.262 | 0.793 | |

**TABLE S6.** *Continuation*

| Group | Response | Predictors | Estimate ± se | | | Statistic | p-value | |
|---|---|---|---|---|---|---|---|---|
| Wasps | Survival | Intercept | -0.762 | ± | 0.142 | 5.370 | <0.001 | *** |
| | | Local temperature | -0.462 | ± | 0.177 | 2.613 | 0.009 | ** |
| | | LST 50 | -0.031 | ± | 0.105 | 0.293 | 0.770 | |
| | | Park area | 0.011 | ± | 0.050 | 0.220 | 0.826 | |
| | | Proximity index | -0.125 | ± | 0.209 | 0.595 | 0.552 | |
| | | NDVI 100 | 0.160 | ± | 0.212 | 0.754 | 0.451 | |
| | | NDVI 800 | 0.000 | ± | 0.033 | 0.009 | 0.993 | |
| | | Grasslands 32 | -0.118 | ± | 0.176 | 0.674 | 0.501 | |
| | | Woody 32 | 0.037 | ± | 0.104 | 0.353 | 0.724 | |
| | | Impervious 32 | -0.124 | ± | 0.184 | 0.676 | 0.499 | |
| | | Plant S | 0.010 | ± | 0.065 | 0.151 | 0.880 | |
| | | Plant H' | -0.011 | ± | 0.073 | 0.149 | 0.881 | |
| | Parasitism | Intercept | -8.288 | ± | 0.288 | 28.759 | <0.001 | *** |
| | | Local temperature | 0.002 | ± | 0.077 | 0.025 | 0.980 | |
| | | LST 800 | 0.052 | ± | 0.141 | 0.372 | 0.710 | |
| | | Park area | 0.006 | ± | 0.030 | 0.184 | 0.854 | |
| | | Proximity index | 0.008 | ± | 0.056 | 0.145 | 0.885 | |
| | | NDVI 100 | -0.019 | ± | 0.093 | 0.206 | 0.837 | |
| | | NDVI 800 | 0.010 | ± | 0.081 | 0.119 | 0.905 | |
| | | Grasslands 32 | 0.014 | ± | 0.096 | 0.147 | 0.883 | |
| | | Woody 32 | 0.014 | ± | 0.097 | 0.140 | 0.889 | |
| | | Impervious 32 | -0.031 | ± | 0.106 | 0.289 | 0.773 | |
| | | Plant S | -0.003 | ± | 0.045 | 0.06 | 0.952 | |
| | | Plant H' | -0.010 | ± | 0.063 | 0.155 | 0.877 | |
| | Females | Intercept | 0.462 | ± | 0.066 | 7.041 | <0.001 | *** |
| | | Local temperature | -0.001 | ± | 0.017 | 0.055 | 0.956 | |
| | | LST 50 | -0.006 | ± | 0.029 | 0.216 | 0.829 | |
| | | Park area | -0.006 | ± | 0.023 | 0.249 | 0.804 | |
| | | Proximity index | -0.013 | ± | 0.069 | 0.190 | 0.849 | |
| | | NDVI 100 | 0.000 | ± | 0.020 | 0.013 | 0.990 | |
| | | NDVI 800 | -0.020 | ± | 0.052 | 0.388 | 0.698 | |
| | | Grasslands 32 | 0.182 | ± | 0.083 | 2.192 | 0.028 | * |
| | | Woody 32 | 0.005 | ± | 0.038 | 0.140 | 0.889 | |
| | | Impervious 32 | -0.015 | ± | 0.046 | 0.330 | 0.741 | |
| | | Plant S | -0.006 | ± | 0.023 | 0.249 | 0.804 | |
| | | Plant H' | -0.105 | ± | 0.107 | 0.980 | 0.327 | |
| | Number of cells | Intercept | 1.169 | ± | 0.038 | 30.920 | <0.001 | *** |
| | | Local temperature | -0.155 | ± | 0.042 | 3.671 | 0.000 | *** |
| | | LST 50 | -0.001 | ± | 0.013 | 0.039 | 0.969 | |
| | | Park area | -0.004 | ± | 0.015 | 0.247 | 0.805 | |
| | | Proximity index | 0.006 | ± | 0.021 | 0.268 | 0.788 | |
| | | NDVI 100 | -0.003 | ± | 0.017 | 0.198 | 0.843 | |
| | | NDVI 800 | -0.001 | ± | 0.011 | 0.061 | 0.951 | |
| | | Grasslands 32 | 0.019 | ± | 0.037 | 0.517 | 0.605 | |
| | | Woody 32 | 0.019 | ± | 0.037 | 0.517 | 0.605 | |
| | | Impervious 32 | -0.013 | ± | 0.031 | 0.424 | 0.672 | |
| | | Plant S | -0.024 | ± | 0.039 | 0.607 | 0.544 | |
| | | Plant H' | -0.027 | ± | 0.044 | 0.619 | 0.536 | |

**TABLE S6.** *Continuation*

| Group | Response | Predictors | Estimate ± se | | | Statistic | p-value | |
|---|---|---|---|---|---|---|---|---|
| Enemy bees | Species richness | Intercept | 0.580 | ± | 0.155 | 3.664 | 0.000 | *** |
| | | Local temperature | 0.000 | ± | 0.037 | 0.002 | 0.999 | |
| | | LST 800 | 0.011 | ± | 0.053 | 0.205 | 0.837 | |
| | | Park area | 0.005 | ± | 0.030 | 0.171 | 0.865 | |
| | | Proximity index | -0.002 | ± | 0.026 | 0.072 | 0.943 | |
| | | NDVI 100 | -0.013 | ± | 0.055 | 0.235 | 0.814 | |
| | | NDVI 800 | 0.003 | ± | 0.035 | 0.096 | 0.924 | |
| | | Grasslands 32 | 0.015 | ± | 0.052 | 0.291 | 0.771 | |
| | | Woody 32 | -0.001 | ± | 0.035 | 0.035 | 0.972 | |
| | | Impervious 32 | -0.007 | ± | 0.042 | 0.169 | 0.866 | |
| | | Plant S | 0.000 | ± | 0.029 | 0.012 | 0.991 | |
| | | Plant H' | 0.020 | ± | 0.063 | 0.314 | 0.754 | |
| | | Host richness | 0.010 | ± | 0.035 | 0.275 | 0.784 | |
| | | Number host nest | 0.008 | ± | 0.003 | 2.819 | 0.005 | ** |
| | Abundance | Intercept | 1.720 | ± | 0.175 | 9.628 | <0.001 | *** |
| | | Local temperature | -0.006 | ± | 0.045 | 0.120 | 0.905 | |
| | | LST 50 | 0.033 | ± | 0.088 | 0.376 | 0.707 | |
| | | Park area | -0.014 | ± | 0.053 | 0.264 | 0.792 | |
| | | Proximity index | 0.001 | ± | 0.022 | 0.038 | 0.970 | |
| | | NDVI 100 | -0.009 | ± | 0.050 | 0.173 | 0.863 | |
| | | NDVI 800 | 0.008 | ± | 0.043 | 0.182 | 0.856 | |
| | | Grasslands 32 | 0.017 | ± | 0.071 | 0.235 | 0.814 | |
| | | Woody 32 | 0.022 | ± | 0.079 | 0.279 | 0.780 | |
| | | Impervious 32 | -0.027 | ± | 0.072 | 0.378 | 0.705 | |
| | | Plant S | -0.025 | ± | 0.066 | 0.374 | 0.708 | |
| | | Plant H' | -0.027 | ± | 0.073 | 0.367 | 0.713 | |
| | | Host richness | 0.000 | ± | 0.024 | 0.009 | 0.993 | |
| | | Number host nest | 0.021 | ± | 0.004 | 4.749 | <0.001 | *** |
| Enemy wasps | Species richness | Intercept | 0.587 | ± | 0.157 | 3.737 | 0.000 | *** |
| | | Local temperature | 0.000 | ± | 0.037 | 0.002 | 0.999 | |
| | | LST 800 | 0.011 | ± | 0.053 | 0.205 | 0.837 | |
| | | Park area | 0.005 | ± | 0.030 | 0.171 | 0.865 | |
| | | Proximity index | -0.002 | ± | 0.026 | 0.072 | 0.943 | |
| | | NDVI 100 | -0.013 | ± | 0.055 | 0.235 | 0.814 | |
| | | NDVI 800 | 0.003 | ± | 0.035 | 0.096 | 0.924 | |
| | | Grasslands 32 | 0.015 | ± | 0.052 | 0.291 | 0.771 | |
| | | Woody 32 | -0.001 | ± | 0.035 | 0.035 | 0.972 | |
| | | Impervious 32 | -0.007 | ± | 0.042 | 0.169 | 0.866 | |
| | | Plant S | 0.000 | ± | 0.029 | 0.012 | 0.991 | |
| | | Plant H' | 0.020 | ± | 0.063 | 0.314 | 0.754 | |
| | | Host richness | 0.010 | ± | 0.035 | 0.275 | 0.784 | |
| | | Host nest abundance | 0.008 | ± | 0.003 | 2.819 | 0.005 | ** |
| | Abundance | Intercept | 0.011 | ± | 0.004 | 3.056 | 0.002 | ** |
| | | LST 800 | 0.239 | ± | 0.220 | 1.085 | 0.278 | |
| | | Park area | -0.005 | ± | 0.031 | 0.163 | 0.871 | |
| | | Proximity index | 0.007 | ± | 0.055 | 0.122 | 0.903 | |
| | | NDVI 100 | 0.015 | ± | 0.073 | 0.208 | 0.835 | |
| | | Grasslands 32 | 0.221 | ± | 0.181 | 1.210 | 0.226 | |
| | | Woody 32 | 0.036 | ± | 0.112 | 0.323 | 0.747 | |
| | | Plant S | -0.003 | ± | 0.028 | 0.124 | 0.901 | |
| | | Host richness | 0.102 | ± | 0.076 | 1.326 | 0.185 | |
| | | Host nest abundance | 0.009 | ± | 0.005 | 1.652 | 0.099 | |

Significance is set at $p < 0.05$. Significance codes are as follows: * $0.05 > p > 0.01$; ** $0.01 > p > 0.001$; *** $p < 0.001$.

S = species richness; H’ = Shannon diversity; NDVI = normalized difference vegetation index; LST = land surface temperature.

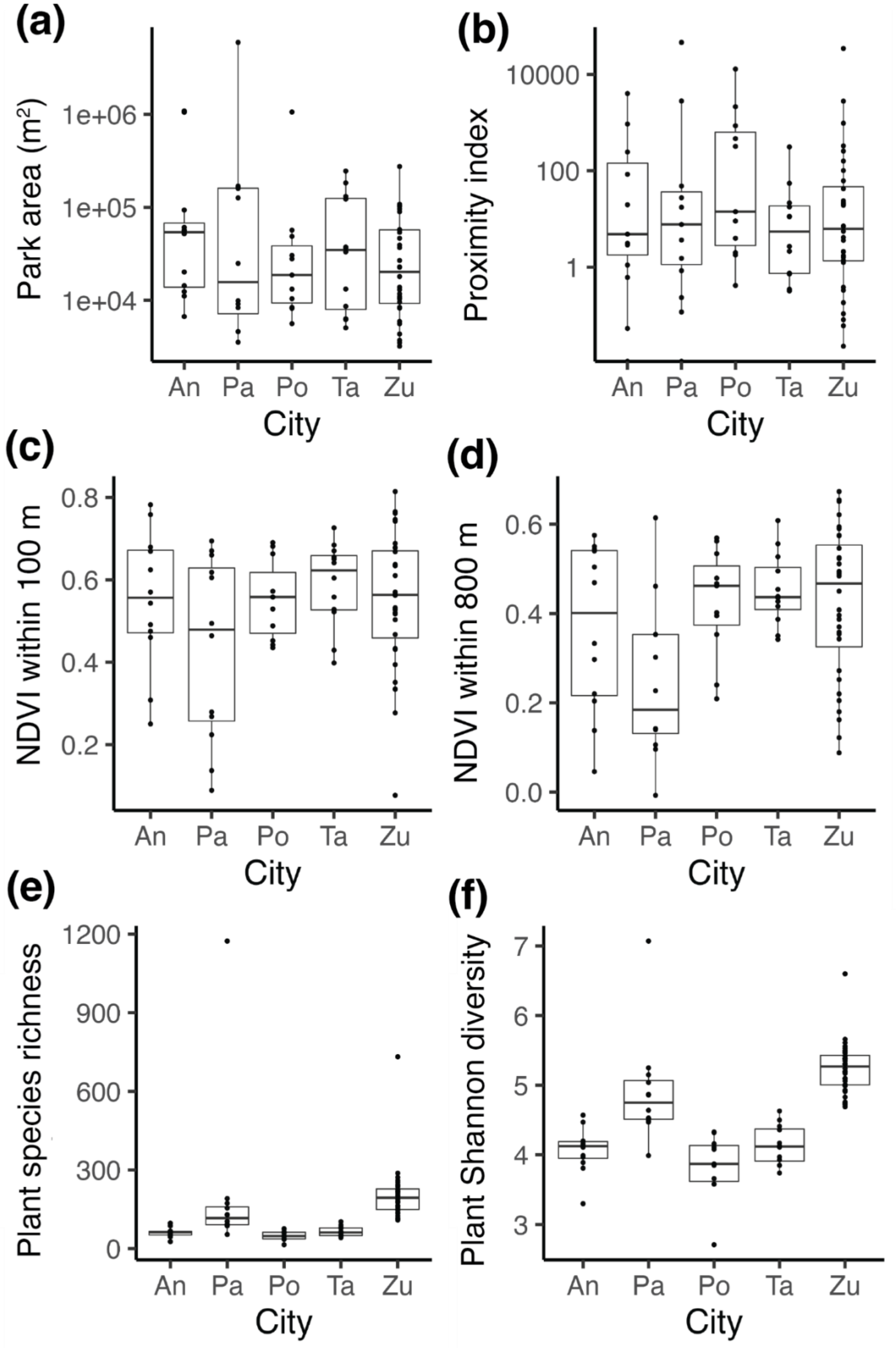


**FIGURE S1.** Boxplots depicting the distribution of values of the predictors across cities. (a) Park area in $m^2$, (b) proximity index, (c) NDVI score within 100 m, (d) NDVI score within 800 m, (e) plant species richness within 100 m, (f) plant Shannon diversity index within 100 m, (g) local temperature at the trap-nest level in °C, (h) LST within 100 m in °C, (i) proportion of artificial surfaces within 32 m, (j) proportion of grasslands within 100 m, and (k) proportion of woody vegetation within 32 m. NDVI = normalized difference vegetation index, LST = land surface temperature. An = Antwerp; Pa = Paris, Po = Poznan; Ta = Tartu; Zu = Zurich.

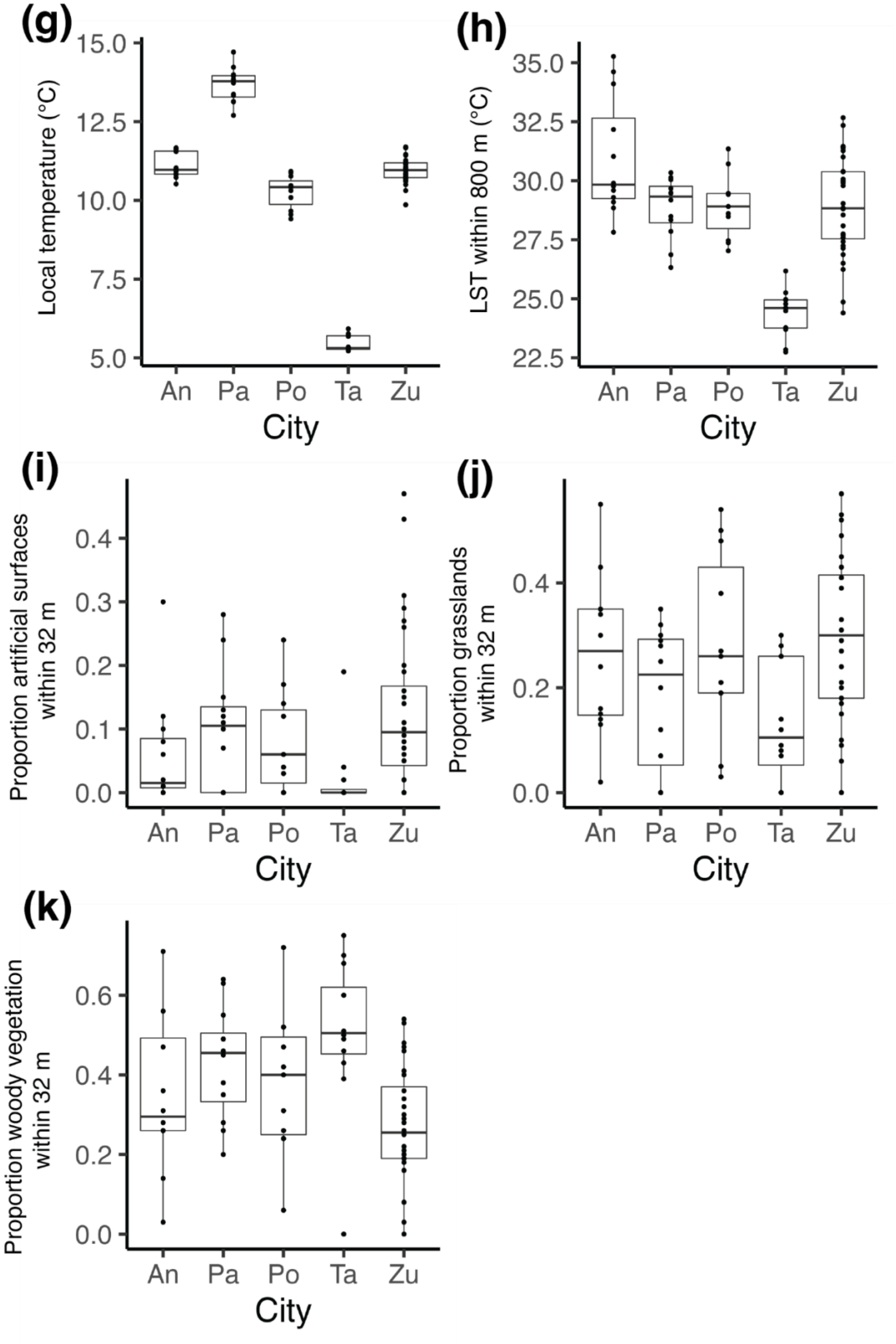


**FIGURE S1.** *Continuation*

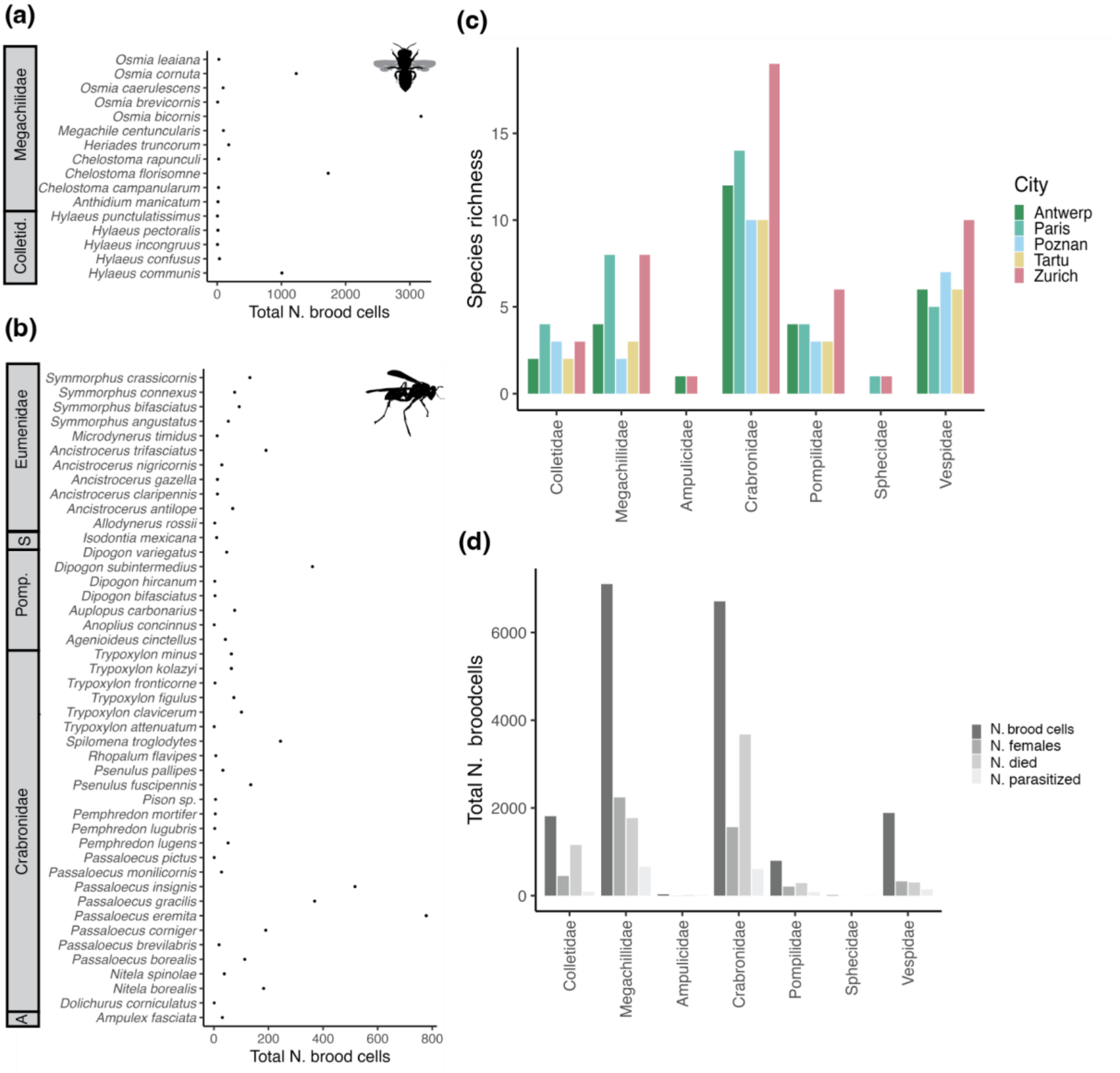


**FIGURE S2.** Community composition. Total number of brood cells recorded per species of cavity-nesting bees (a) and wasps (b). Species are sorted according to their families. Morphospecies (individuals identified at the genus or family level) are not shown. Note that *Pison* sp. has been included here as it is the only record of the genus. Species richness (c) and total number of brood cells (d) in the five studied cities of the different host families of cavity-nesting bees (i.e. Colletidae and Megachilidae) and wasps (i.e. Ampulicidae, Crabronidae, Pompilidae, Sphecidae, Vespidae). Note that Zurich has 32 study sites, whereas the remaining four cities (Antwerp, Paris, Poznan and Tartu) each have 12 study sites. A = Ampulicidae; Colletid = Colletidae; S = Spechidae; Pomp = Pompilidae.

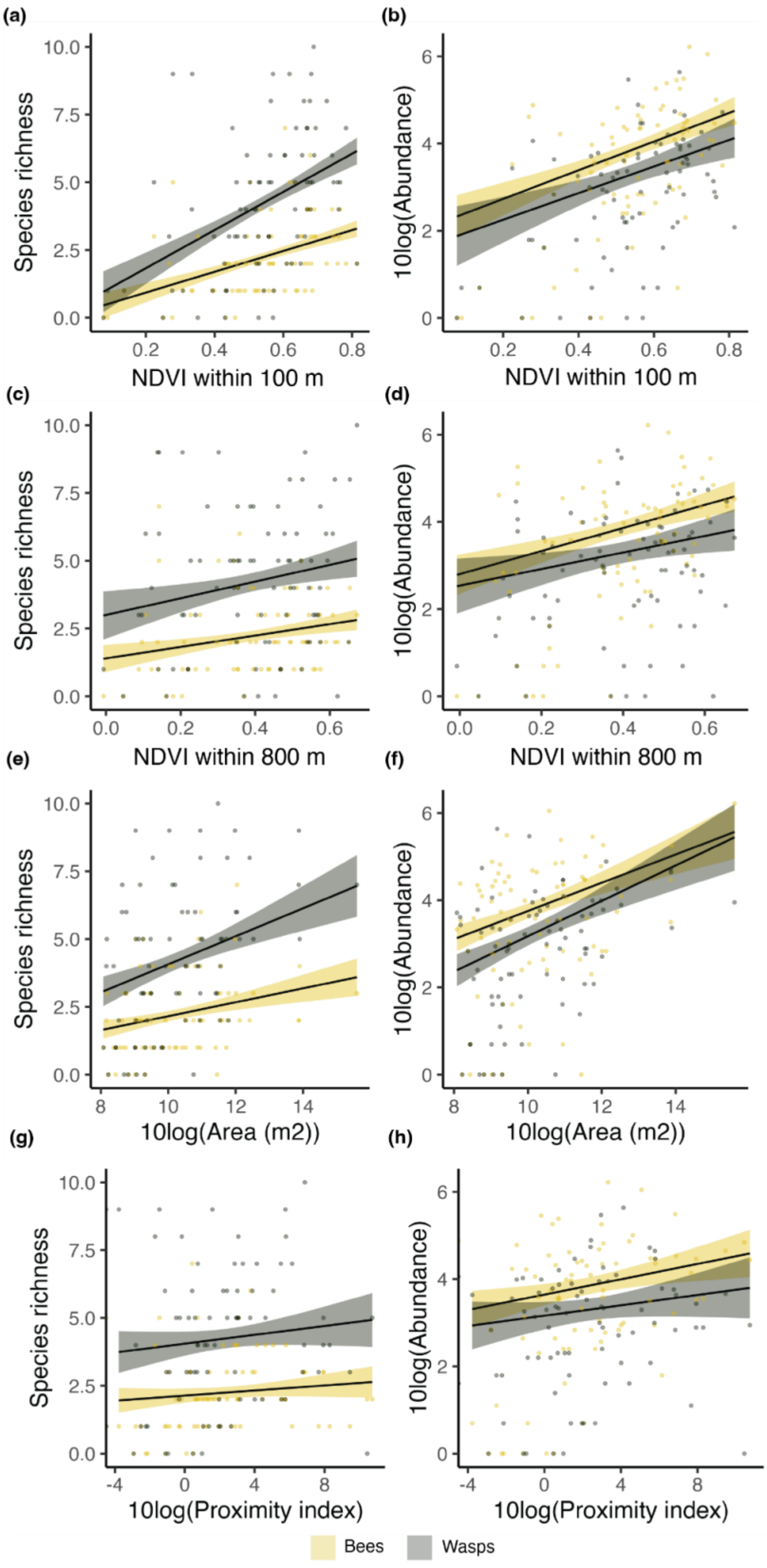


**FIGURE S3.** Generalized linear models (GLMMs) depicting the relationship between the predicted species richness or the predicted abundance of hatched offspring of cavity-nesting bees and wasps and the proxies for habitat amount (a–h), temperature (i–l, see figure continuation below), resource availability (m–v), and top-down controls (w–x). Points show the raw data and solid lines show the predicted values obtained using averaged GLMMs. Coloured bands indicate the 95% confidence intervals. Model results are shown in Table S6. Plant S = Plant species richness; Plant H’ = Plant Shannon diversity.

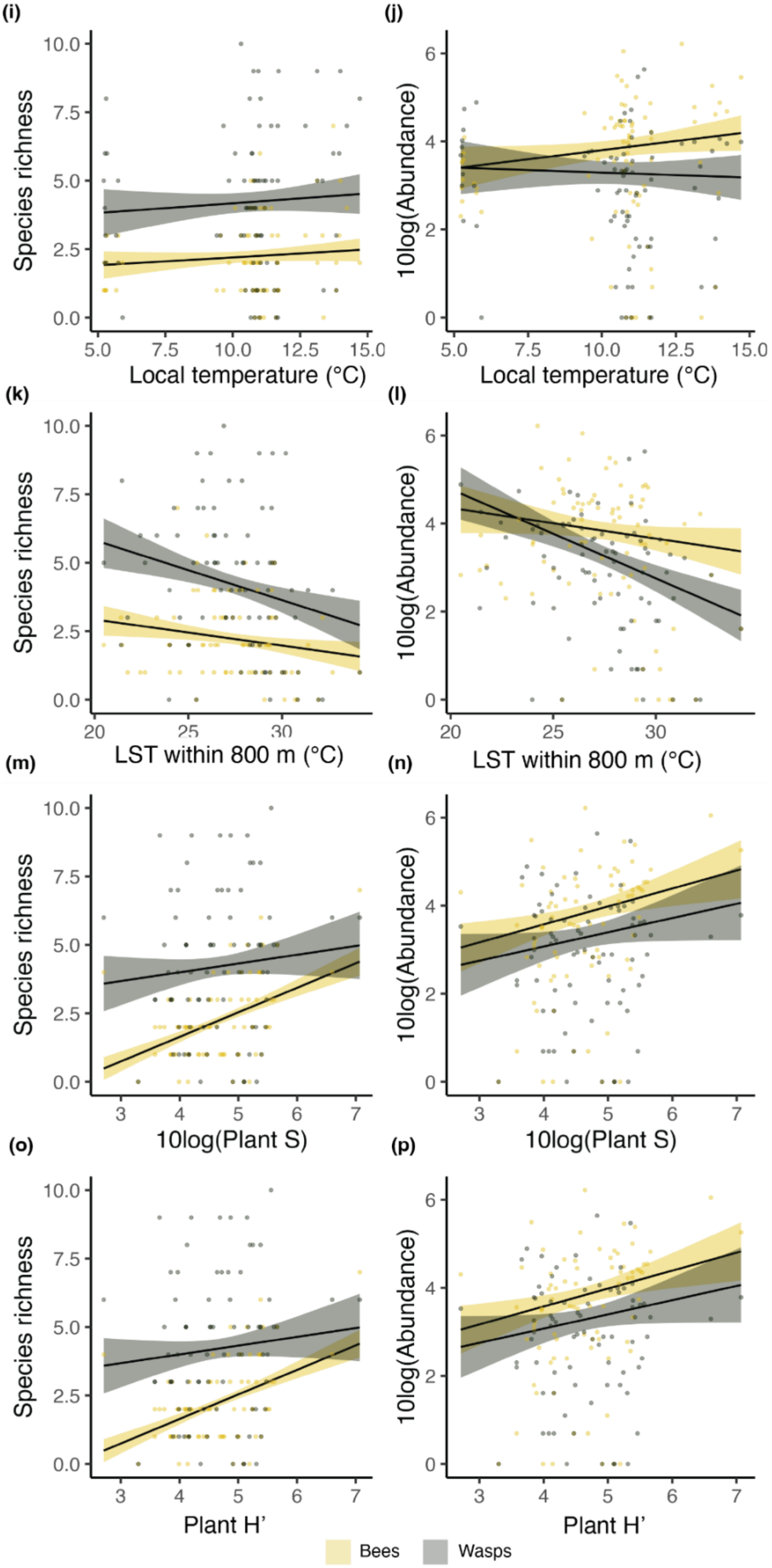


**FIGURE S3.** *Continuation.*

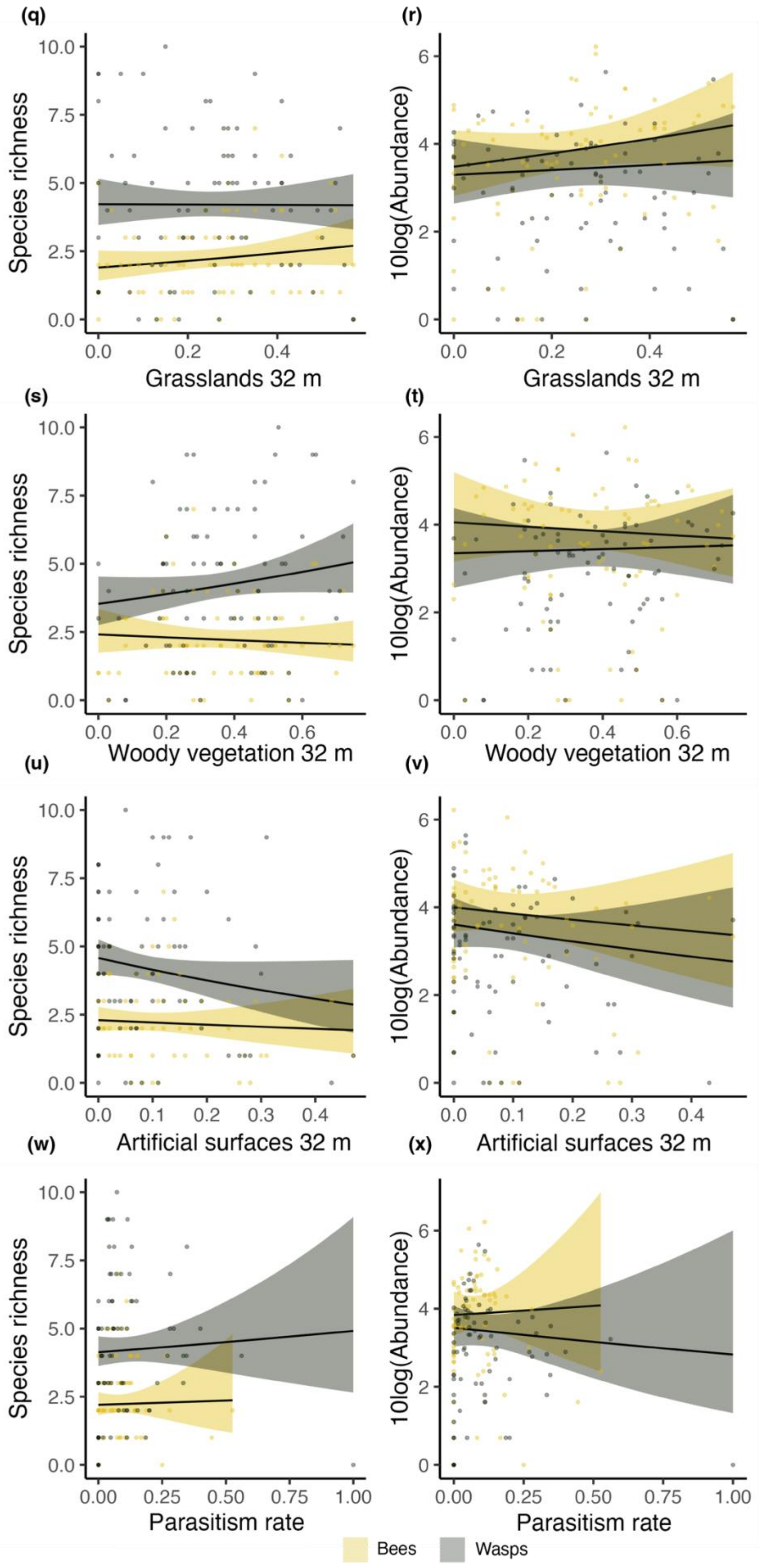


**FIGURE S3.** *Continuation.*

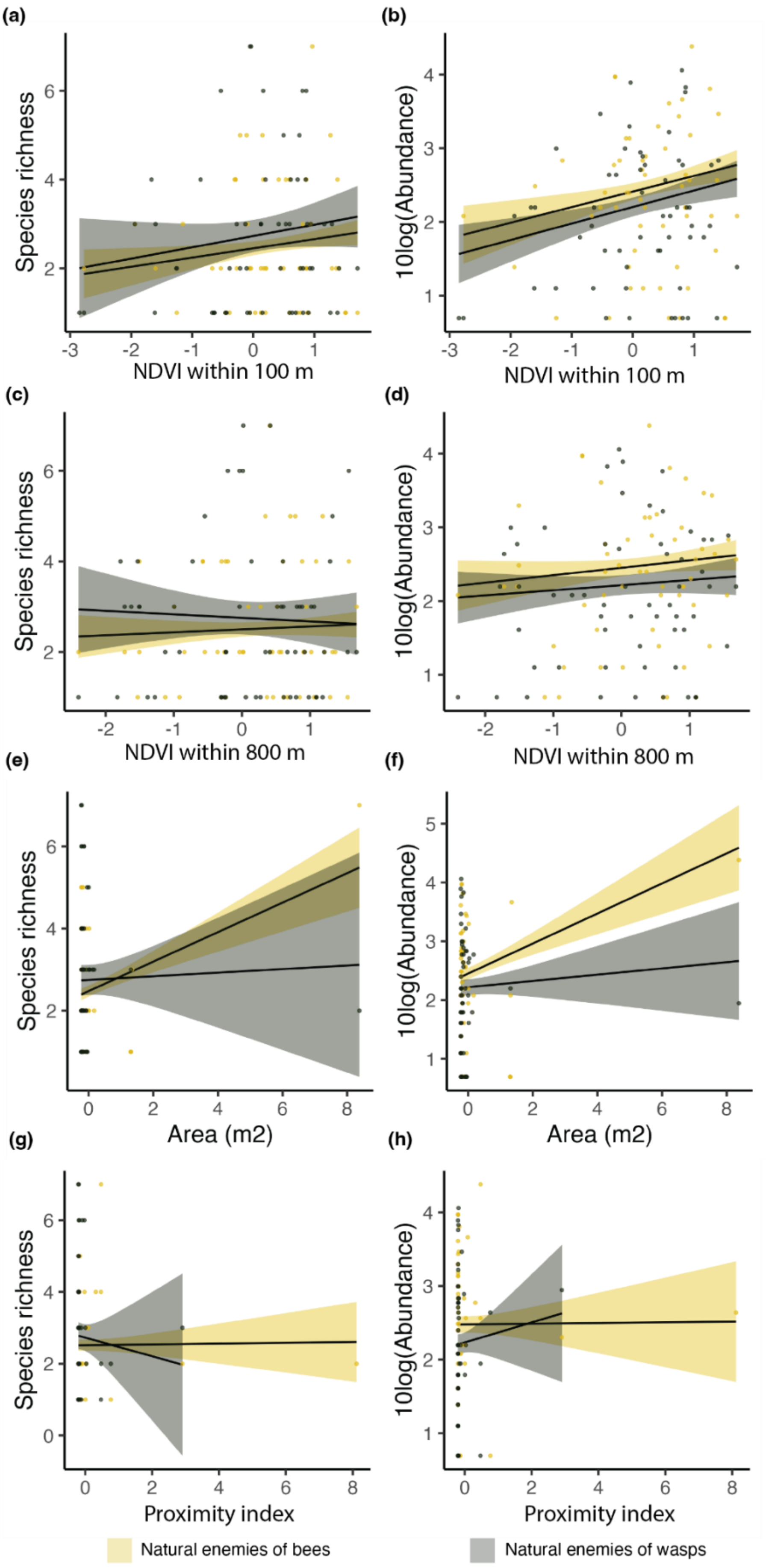


**FIGURE S4.** Generalized linear models (GLMMs) depicting the relationship between the predicted species richness or the predicted abundance of the natural enemies of cavity-nesting bees and wasps and the proxies for habitat amount (a–h), temperature (i–l, see figure continuation below), resource availability (m–v), and top-down controls (w–x). Points show the raw data and solid lines show the predicted values obtained using averaged GLMMs. Coloured bands indicate the 95% confidence intervals. Model results are shown in Table S6.

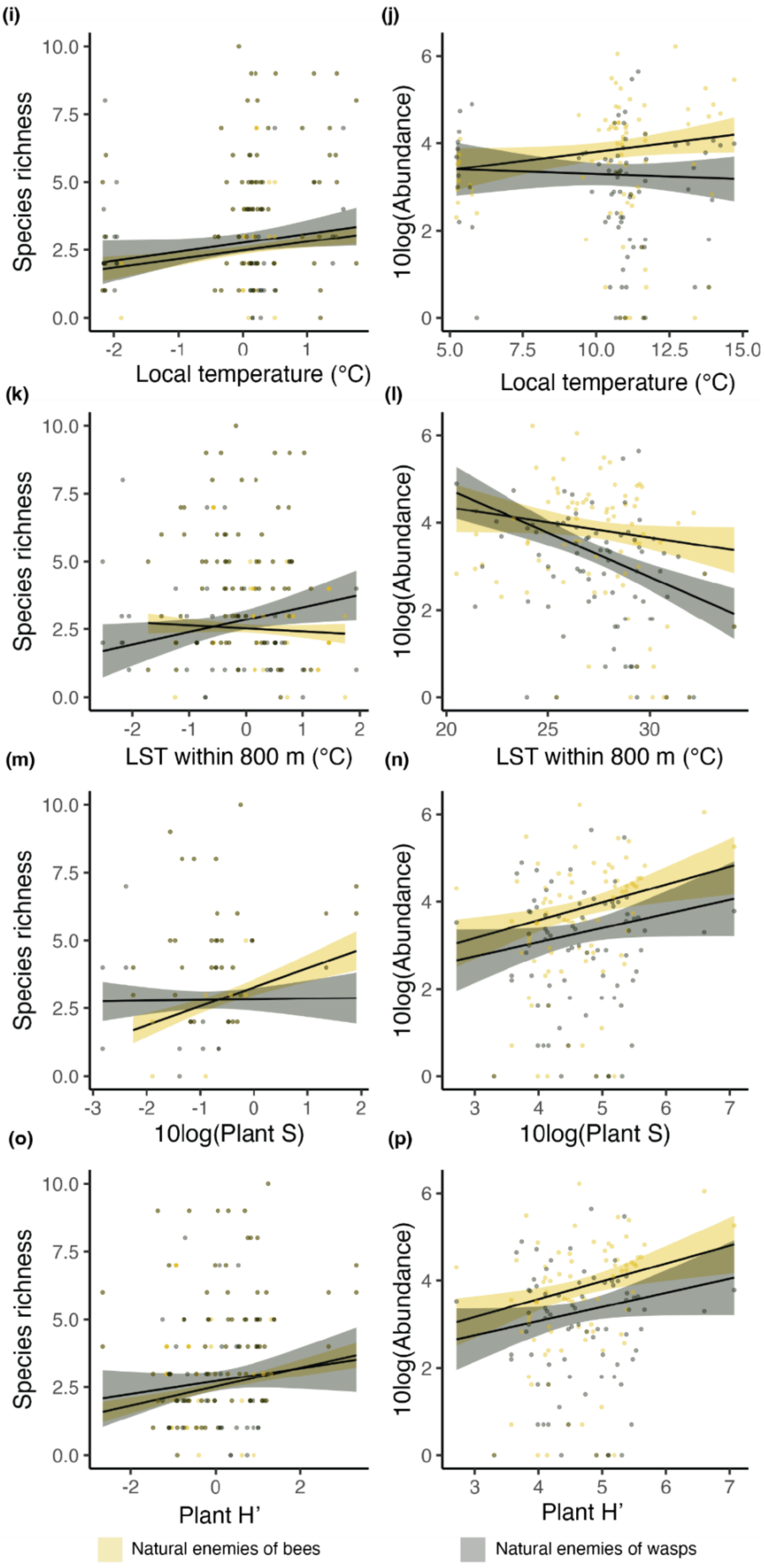


**FIGURE S4.** *Continuation.*

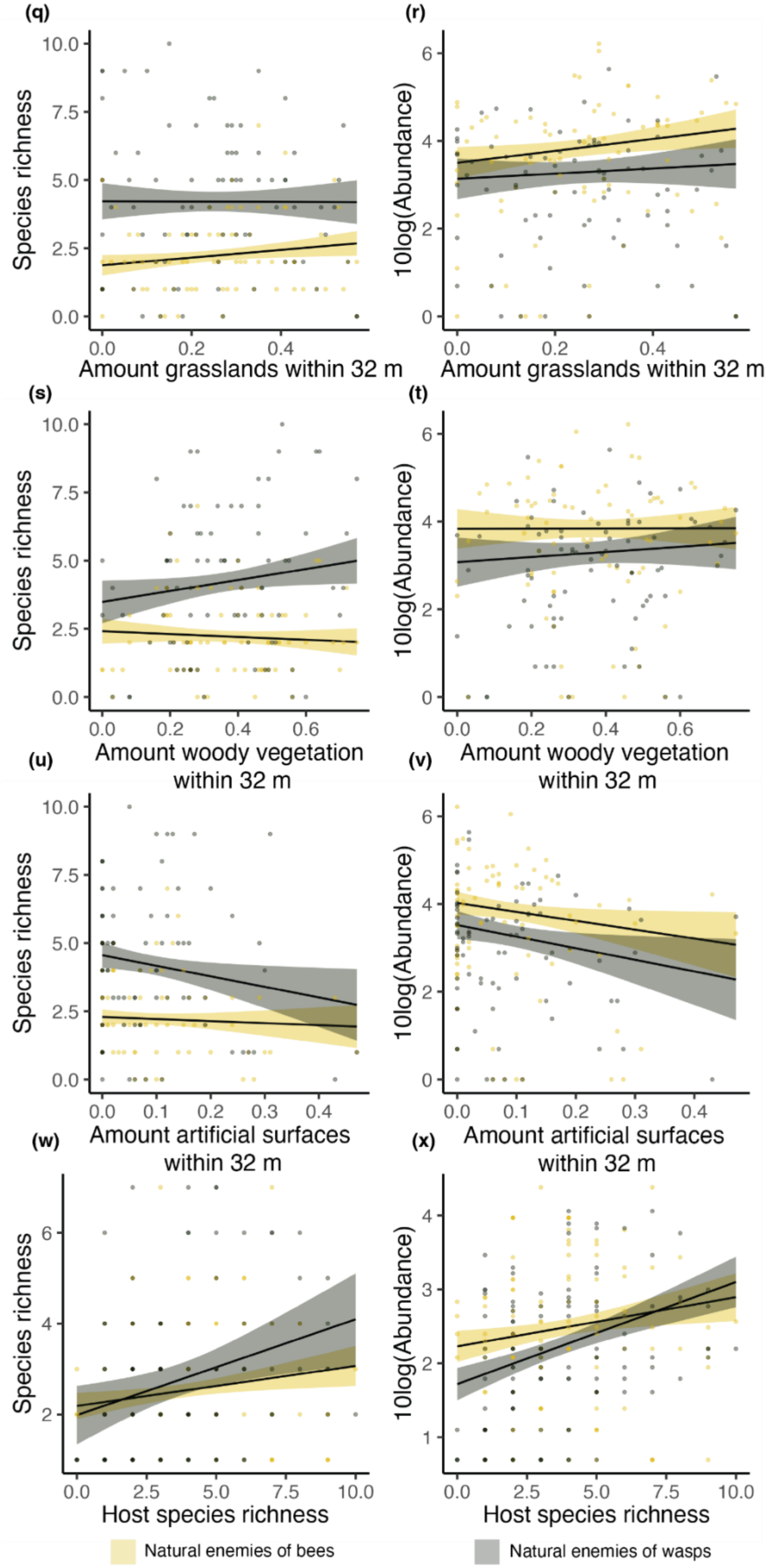


**FIGURE S4.** *Continuation.*

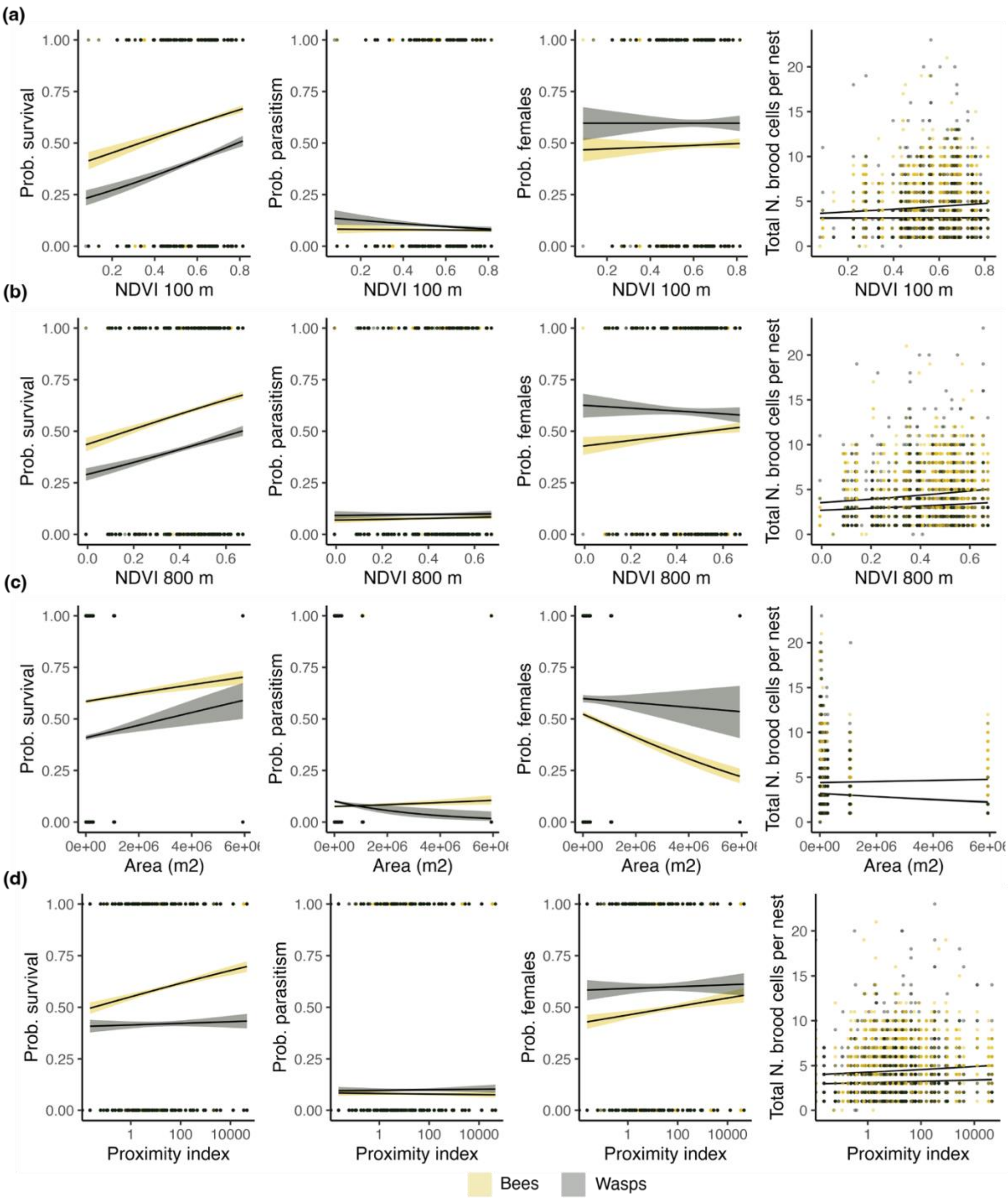


**FIGURE S5.** Generalized linear models (GLMMs) depicting the relationship between the predicted probability of survival, probability of parasitism, probability of female, or the total number of cells for both cavity-nesting bees and wasps and the proxies for habitat amount (a–d, g–k) and temperature (e–f). Points show the raw data and solid lines show the predicted values obtained using averaged GLMMs. Coloured bands indicate the 95% confidence intervals. Model results are shown in Table S6.

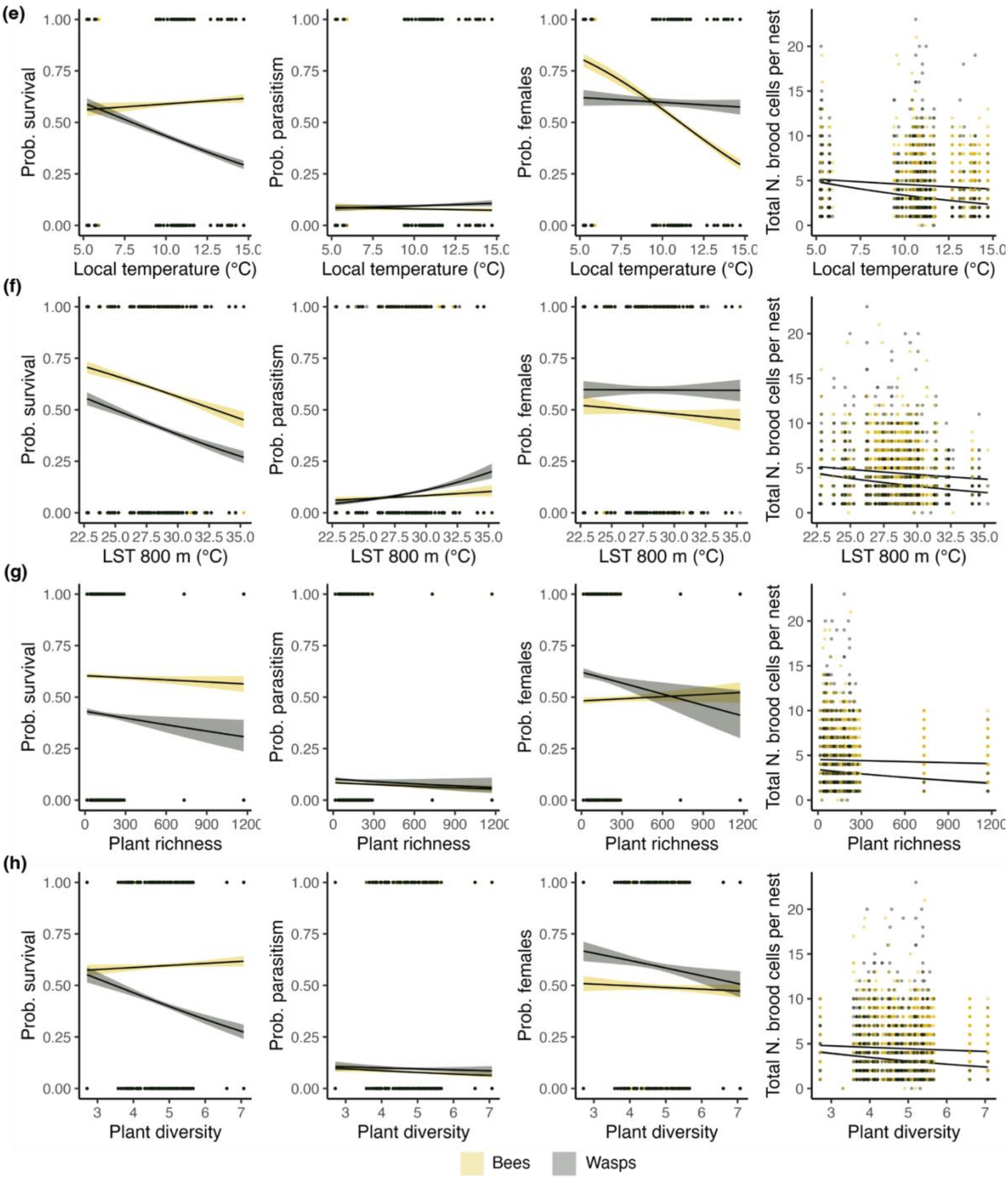


**FIGURE S5.** *Continuation.*

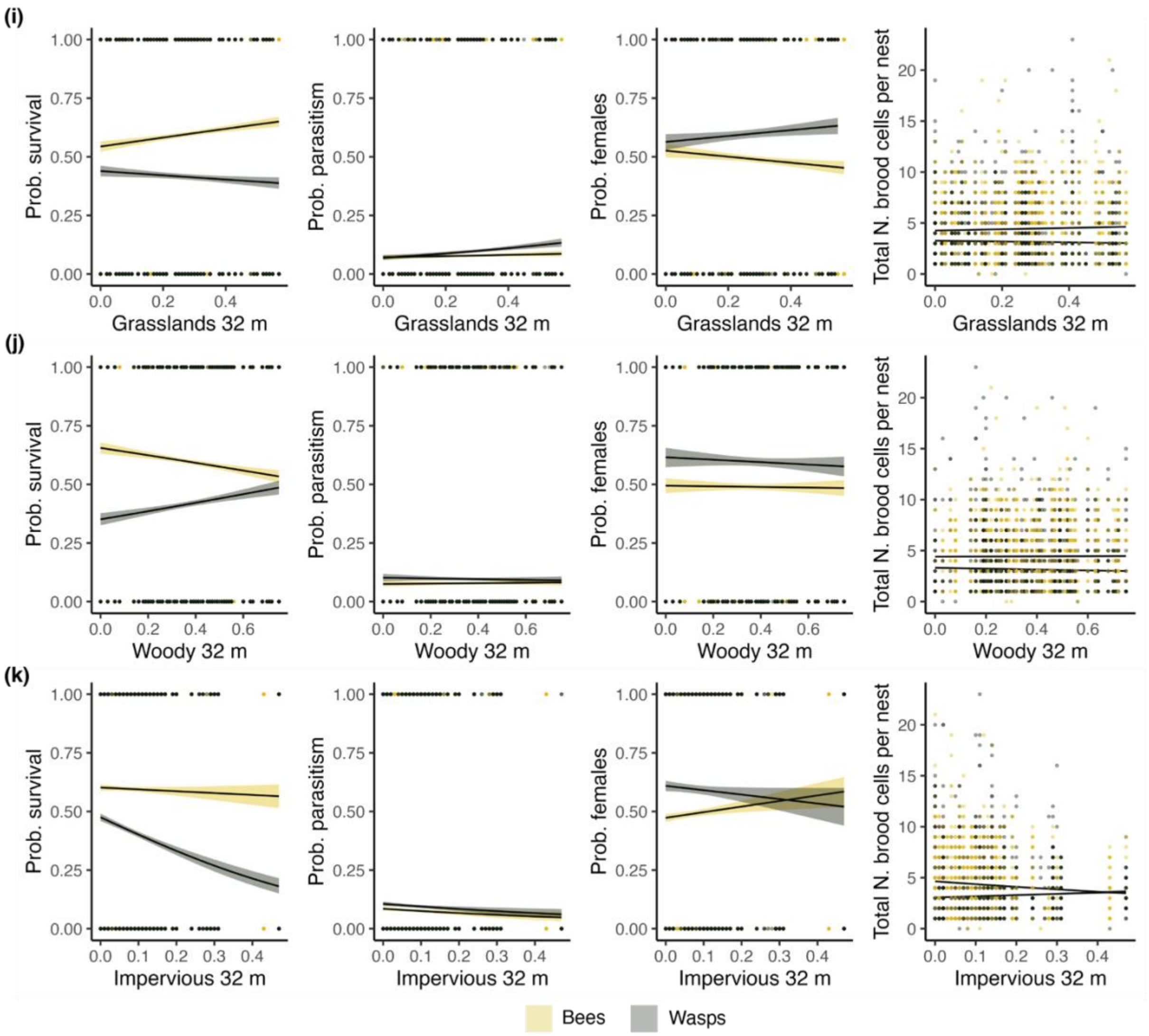


**FIGURE S5.** *Continuation.*
